# Automating scientific annotations for open transcriptomic profiles via multi-stage agents

**DOI:** 10.64898/2026.08.19.745739

**Authors:** Xiaodan Zhang, Shreya Paithankar, Jing Pu, Mehtab Sardar Murtaza, Rama Shankar, Dmitry Leshchiner, Shubham Koirala, Zsolt Palmer, Rance Nault, Xiaopeng Li, Yuying Xie, Bin Chen

## Abstract

Public transcriptomic repositories contain millions of samples, yet their large-scale reuse is hindered by heterogeneous and inconsistently reported metadata. In the Gene Expression Omnibus (GEO), key biological information is often distributed across study- and sample-level records, requiring context-dependent interpretation. Here we present GEOMeta, a large language model (LLM)-based multi-stage workflow with task-specialized agents for automated GEO metadata curation. The pipeline separates metadata retrieval, task-specific information extraction, field standardization, ontology mapping and quality control. Using GEOMeta, we generated standardized annotations for approximately 600,000 human bulk RNA-seq samples. To demonstrate its utility, we benchmarked transcriptome representation models for predicting sex, age, tissue and disease from transcriptome embeddings. We further prospectively annotated newly submitted GEO studies and evaluated 22 frontier LLMs. Recent open-source Flash models achieved annotation quality comparable to leading reasoning models while reducing costs by an order of magnitude. GEOMeta provides a scalable resource and reproducible framework for metadata curation.

## Introduction

Large, diverse, and high-quality datasets have repeatedly driven major advances in machine learning (ML) and deep learning (DL). Outstanding examples include ImageNet ^1,2^, which revolutionized computer vision and catalyzed the development of deep neural networks and convolutional neural networks, and Protein Data Bank for experimentally determined 3D protein structures, which enabled the development of the AlphaFold family of models that revolutionized the learning of protein structure ^3,4^. Transcriptomics is among the most widely used molecular profiling modalities in biomedical research. Analogously, large collections of transcriptomic profiles hold the promise of enabling the learning of generalizable representations of cellular and disease states and uncover new biology. Several models have been developed for unsupervised representation learning of bulk transcriptomic data ^5–9^, but many have been trained predominantly on cancer-focused resources, particularly The Cancer Genome Atlas (TCGA), potentially limiting their generalizability across tissues, diseases and experimental settings. Foundation models for single-cell transcriptomics have recently shown promise across a range of downstream tasks ^10–13^, but their performance is similarly dependent on the diversity, quality and annotation of the data used for training and evaluation.

To overcome these limitations and train generalizable models, researchers require access to massive, highly diverse datasets. The Gene Expression Omnibus (GEO), established by the National Center for Biotechnology Information (NCBI), is the largest publicly accessible repository for transcriptomics data. GEO contains millions of samples from hundreds of thousands of studies and continues to grow rapidly. Reuse of open transcriptome profiles has supported disease signature metanalysis ^14,15^, drug repurposing ^16,17^, biomarker discoveries ^18,19^, and preclinical model evaluation ^20,21^. In addition, resources such as ARCHS4 ^22^, Recount2 ^23^, and Treehouse ^24^ have reprocessed and normalized GEO transcriptome profiles to support large-scale downstream analysis. Nevertheless, as those resources lack structured meta data, most applications still rely on relatively small, manually curated subsets.

Standardized sample-level metadata are therefore essential for fully exploiting harmonized transcriptomic profiles and supporting representation learning ^25–27^. Although GEO provides structured submission fields, many fields are optional, and biologically relevant information is frequently distributed across study summaries, experimental-design descriptions and sample characteristics. Equivalent concepts may be described using different field names or values, for example, “strain” versus “Strain background,” or “bleomycin treated” versus “treatment: BLM.” Conversely, similar terms can have different meanings depending on the study context. A compound name may denote the principal perturbation in one study but a background treatment, culture condition or comparator in another. Correct annotation can therefore require joint interpretation of study-level and sample-level records rather than extraction of isolated phrases.

Early efforts to annotate public transcriptomic metadata relied largely on manual curation and were typically restricted to selected organisms, assays, scientific questions, or biological domains ^27–31^. Subsequent approaches used ontologies, rule-based systems, named-entity recognition and supervised language models to normalize sample descriptions and infer missing annotations ^26,32,33^. These methods substantially improved metadata accessibility but remain challenged by ambiguous terminology, incomplete reporting and annotations that require contextual or cross-field reasoning. Single-pass prompting of a general-purpose language model presents similar limitations ^34^: extraction, semantic interpretation, normalization, controlled derivation and ontology mapping are distinct tasks, and errors introduced at one stage can propagate into subsequent annotations.

Recent large language models (LLMs) can process longer textual contexts and perform flexible, task-specific reasoning ^35–37^, creating an opportunity to automate scientific metadata curation at greater scale. However, reliable curation requires more than unconstrained text generation. Annotations must conform to predefined schemas, distinguish explicit evidence from inferred attributes, preserve provenance and remain robust across diverse experimental scenarios. LLM outputs may also vary across prompts and models or contain hallucinated results^38^. The utility of LLMs for large-scale scientific curation therefore depends on workflow design, programmatic constraints and systematic field-level evaluation.

Here we present GEOMeta, an automated framework for curating GEO metadata through staged, role-specific LLM processing. The framework separates contextual extraction, validation, field-level standardization, controlled derivation and ontology mapping into modular and independently reviewable components. These modules operate under predefined schemas and are coupled to programmatic validation and quality-control procedures, enabling consistent processing of heterogeneous GEO records while preserving intermediate outputs and supporting error tracing. The implementation uses a configurable LLM interface, allowing the annotation backend to be specified through the model name, application programming interface (API) endpoint and authentication credentials.

We applied GEOMeta to human bulk RNA-sequencing studies and generated standardized sample-level annotations for approximately 600,000 samples, including a subset linked to processed expression profiles. We evaluated annotation quality during pipeline development using a manually curated 1,000-sample development and benchmarking set, text-evidence validation, and comparisons with external curated resources and expert review of a separate cohort of recently submitted GEO studies. We further assessed alternative extraction strategies and LLM backends, with particular attention to context-dependent fields including disease, tissue and perturbation. To evaluate the downstream utility of the resulting resource, we benchmarked multiple state-of-the-art transcriptomic representation-learning methods using study-aware data partitions and prediction tasks for sex, age, tissue and disease.

### Overview of the Automated Annotation Framework

We developed GEOMeta to automate the curation of GEO metadata by combining the natural-language processing and reasoning capabilities of LLMs with minimal manual intervention. The resulting annotations are structured, machine-readable and were designed to support the FAIR principles of findability, accessibility, interoperability and reusability ^39^ (Fig. 1A and Methods). We applied the framework to human bulk RNA-sequencing samples and, for the expression-matched subset, linked the resulting annotations to processed gene-expression profiles available through ARCHS4 ^39–41^. Guided by downstream analytical applications, we curated RNA library type, RNA source, experimental setting, study-level perturbation, sample-level perturbation, disease, tissue, age (age group) and sex. Given the growing demand for diverse chemical perturbation-induced transcriptomic profiles in drug discovery ^39–41^, we further focused on chemically perturbed samples and annotated their cellular context, treatment duration and dose. For this subset, the standardized metadata were linked to the corresponding expression profiles to support large-scale data mining, transcriptomic representation learning and the development and validation of patient-level inference models. We subsequently demonstrated the utility of GEOMeta by benchmarking multiple state-of-the-art representation-learning methods on prediction tasks for sex, age, tissue and disease and released the resulting annotations, predefined data splits, and trained baseline models as community resources.

**Figure 1.**
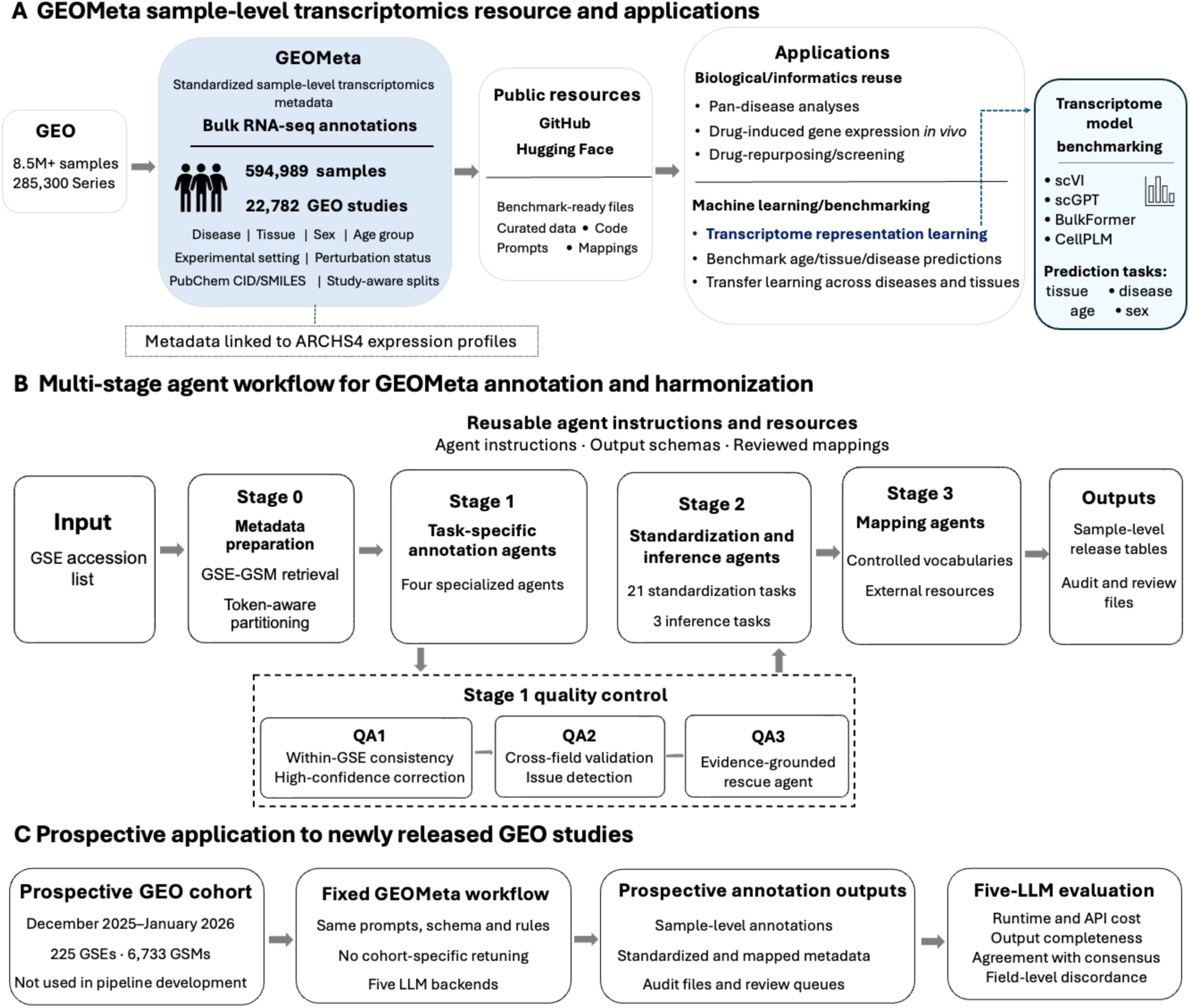
Automated annotation framework for GEO metadata through a multi-stage agent workflow. **A**. GEOMeta resource overview and downstream applications. **B**. Multi-stage agent workflow for GEOMeta annotation and harmonization. **C**. Prospective application to GEO studies submitted between December 2025 and January 2026, a period excluded from the initial GEOMeta release.

However, achieving reliable annotation at this scale required addressing the heterogeneity and complexity of GEO records, which cannot be adequately handled using a single prompt, even with advanced LLMs. Biologically relevant information is frequently distributed across study- and sample-level records, described ambiguously or represented inconsistently among studies. In addition, extraction, semantic interpretation, normalization, controlled derivation and ontology mapping constitute distinct tasks that require different forms of reasoning and validation. We therefore designed a multi-stage agent workflow that decomposes metadata curation into modular and independently reviewable steps (Fig. 1B and Methods). The framework combines task-specific LLM agents for context-aware extraction, field-level standardization, ontology-guided refinement, and rule-based post-processing, reducing the propagation of ambiguous or inconsistent intermediate outputs. GEOMeta used a curated library of 32 task-specific prompt templates spanning 2,469 non-empty lines and 28,348 words: four task-specific annotation agents, 21 field-specific standardization agents, three controlled inference agents, and four controlled-vocabulary and external-resource mapping agents. To improve interoperability, selected entities were further mapped to widely used standardized representations, including SMILES strings for compounds and Medical Subject Headings (MeSH) identifiers for diseases.

To evaluate the feasibility of prospective deployment, we applied the pipeline to more than 5,000 samples from 225 GEO Series records submitted between December 2025 and January 2026, a temporal hold-out period excluded from the initial GEOMeta release (Fig. 1C). These studies were selected from recently submitted GSEs with processed expression profiles available in the recent release of ARCHS4. The framework was initially developed using models from the GPT family, primarily GPT-4, which represented the state of the art during development. Given the rapid evolution of LLMs, we next assessed whether contemporary models can support automated scientific curation. Because LLM performance, computational cost and inference speed vary substantially among models, we evaluated 22 widely used models available through OpenRouter using an internal benchmark cohort comprising five GEO Series submitted by our laboratory. Based on output completeness, consensus concordance, cost and processing efficiency, we selected five models to comprise the final annotation pipeline for the annotation of 5,000 prospective samples. Each model annotated the prospective cohort independently and field-level majority voting was used to construct a consensus dataset for downstream evaluation. The resulting prospectively annotated dataset was then used in downstream evaluations of transcriptomic representation-learning methods.

## Results

### The GEOMeta Dataset

The final GEOMeta release contains 594,989 curated GSM-GSE records, corresponding to 594,304 unique GSM accessions from 22,782 GSEs. Study sizes varied substantially, with a median of 12 samples per study. Coverage was nearly complete for experimental setting (100.0%), study-level perturbation (100.0%), RNA library type (99.96%) and sample-level perturbation (99.17%). RNA source was available for 592,201 samples (99.5%). Disease and tissue were annotated for 98.5% and 87.4% of samples, respectively. The demographic attributes were less frequently reported: sex was available for 31.8%, age group for 16.2% and exact age for 15.7% of samples. For chemical perturbation (CP) samples, dosage and duration were reported for 114,094 samples (19.2%) and 157,444 samples (26.5%), respectively (Supplementary Table 1).

Standardization substantially reduced metadata fragmentation (Fig. 2A). Six sex-label variants were unified into two standardized values. Age labels decreased from 5,253 to 4,170 standardized values after unit and format harmonization. Numeric ages were mapped to 12 predefined age groups, whereas broad non-numeric labels, custom age intervals and developmental-age expressions were retained as reported, resulting in 55 distinct final age-group values. Tissue labels showed the greatest compression, decreasing from 2,246 terms to 56 categories aligned to the Human Protein Atlas ^42^ after resolving synonyms and composite anatomical descriptions. Disease labels were consolidated from 3,454 terms to 1,235 standardized terms through synonym resolution and alignment to the Comparative Toxicogenomics Database (CTD) ^43^ (Fig. 2A).

**Figure 2.**
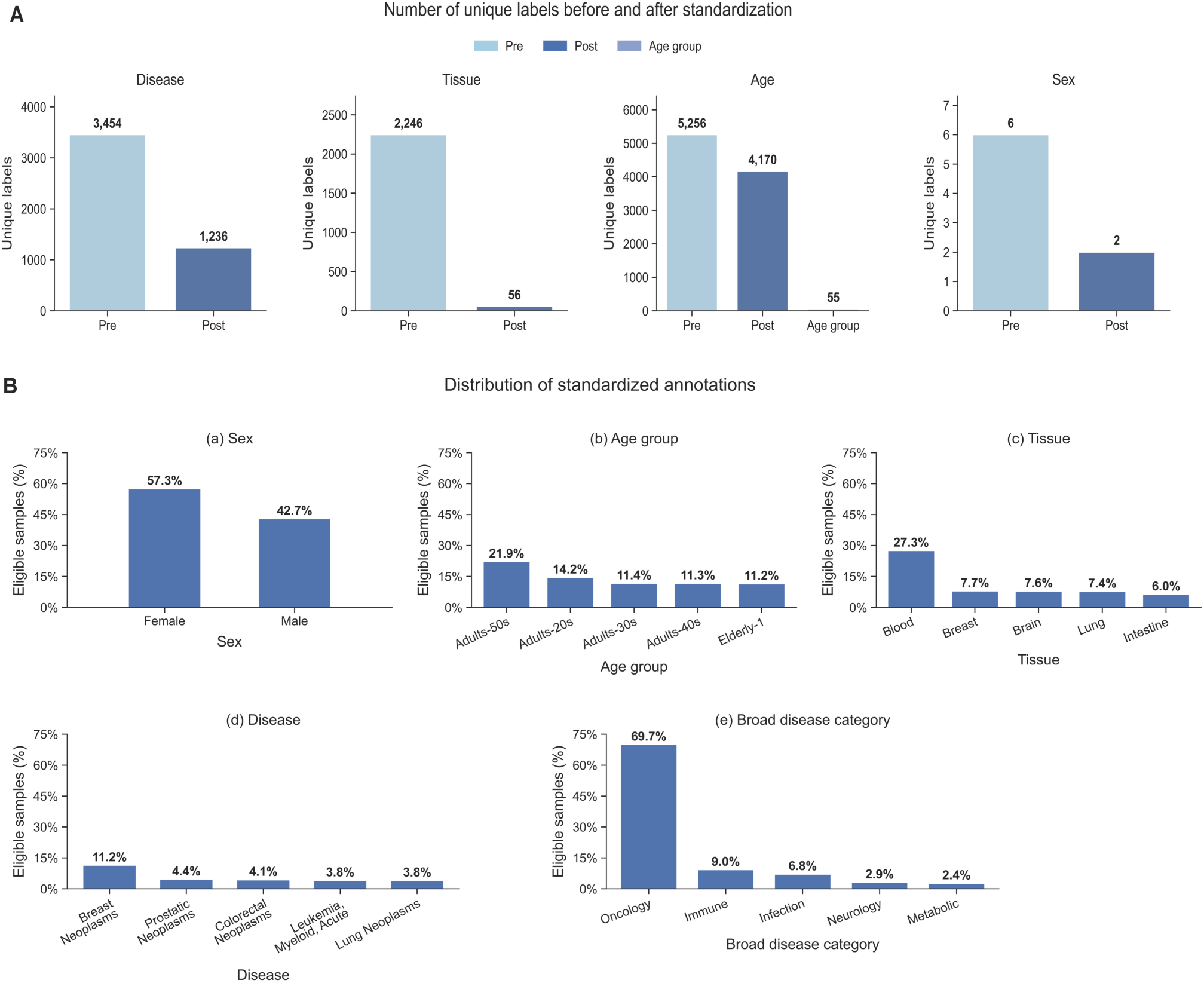
GEOMeta sample summary. **A**: Number of unique labels before and after standardization across five key metadata categories. The panels illustrate vocabulary compression achieved by ontology-aligned normalization and rule-based post-processing. **B**: Distribution of standardized experimental, demographic, and biological annotations in the GEOMeta dataset. Only these fields were used for downstream representation learning.

GEOMeta spans diverse biological and experimental contexts (Fig. 2B). Among samples with reported sex, 57.3% were female and 42.7% were male. Age-group annotations were dominated by adults in their 20s to 50s, whereas older adults, adolescents, children and infants were less frequently represented. Blood was the most represented tissue (27.3%), followed by breast (7.7%), brain (7.6%), lung (7.4%) and intestine (6.0%); the remaining 44.0% covered a broad range of organs, including skin, liver, kidney and reproductive tissues.

Disease-associated annotations were assigned to 317,012 samples (53.3%), whereas 268,732 samples (45.2%) were classified as normal or control related and 9,245 samples (1.6%) lacked a disease label. Disease-associated samples encompassed 1,232 standardized terms with 1,154 unique disease identifiers. Breast neoplasms were the most frequently represented individual disease (11.2% of disease-associated samples), followed by prostatic neoplasms (4.4%), colorectal neoplasms (4.1%), acute myeloid leukemia (3.8%) and lung neoplasms (3.8%). The standardized terms were further organized into 24 broad disease categories. Among the 315,163 samples assigned a broad disease category, oncology was the most frequent category (69.6%), followed by immune disorders (9.0%), infectious diseases (6.8%), neurological disorders (2.9%), metabolic diseases (2.4%) and genetic disorders (2.4%).

Chemical perturbations were also standardized and linked to PubChem Compound ID^44^. After excluding composite treatments, 66.7% of eligible single-compound terms were successfully mapped to PubChem. These mappings represented 72,117 compound-treated samples and 2,787 unique PubChem compounds.

### Multi-pronged Evaluation of the Annotation Agent Pipeline

We developed a multi-pronged evaluation strategy to assess the annotation pipeline before generating the final data release. Because comprehensive ground-truth annotations were not available for all metadata fields, we first assembled a development set of 1,000 representative samples spanning diverse experimental and reporting scenarios. Initial annotations were generated using GPT-4 and subsequently reviewed and corrected manually. This dataset was used exclusively for agent-instruction development, workflow optimization and internal comparison of pipeline iterations. Following iterative prompt refinement and ablation testing, the pipeline achieved consistent agreement with the reviewed development annotations across the evaluated metadata fields. Agreement with expert-reviewed annotations reached 96.8% for sex, 92.8% for experimental setting, 92.3% for age, 95.1% for disease and 79.3% for tissue. Because the same benchmark was used during prompt refinement, these values reflect internal agreement on the development benchmark rather than performance on an independent test set. We then applied the optimized pipeline to approximately 600,000 samples and assessed annotation consistency against corresponding fields available in independently published databases containing manually curated GEO metadata. For attributes that can be inferred from transcriptomic profiles, we additionally trained predictive models to identify potentially discordant annotations. This model-in-the-loop strategy enabled targeted review of suspicious records and, in some cases, revealed inconsistencies between the curated annotation and the original GEO submission. Finally, we deployed the pipeline for evaluation by domain experts using datasets relevant to their interest.

Prompt design affected annotation availability and cross-prompt agreement differently across metadata fields. Across the 1,000-sample prompt-comparison dataset, Stage 1 outputs differed between the task-specific agent configuration and the generic structured configuration for 50.1% of tissue and 52.0% of perturbation annotations. After identical downstream standardization and mapping, final outputs differed in 37.3% and 53.4% of samples, respectively. For tissue, this difference was driven mainly by annotation availability: missingness was 20.6% with task-specific prompts and 53.8% with the generic prompt, whereas only five shared outputs were discordant, yielding 98.9% concordance. Perturbation missingness was likewise lower with task-specific prompts (10.0% versus 31.8%), but 276 shared outputs remained discordant, yielding 58.3% concordance. Disease outputs differed in 14.1% of samples at Stage 1 and 7.8% after mapping, with 99.1% concordance among shared final outputs. RNA-source outputs showed low cross-prompt concordance (12.4%). RNA library and experimental setting showed more modest differences, whereas age group and sex were fully concordant among shared final outputs (Supplementary Table S2).

To further assess annotation consistency, we compared GEOMeta with two independently curated external resources. For disease and tissue annotations, we aligned overlapping samples with DiSignAtlas ^28^, a manually curated database focused on disease-associated transcriptomic signatures. Among the 19,879 samples matched between GEOMeta and DiSignAtlas,13,075 were disease-associated samples with specific disease labels. For these samples, GEOMeta disease annotations were concordant with DiSignAtlas for 96.59% when exact, synonymous and broader-category matches were included, whereas 3.41% remained unmatched. Across 19,879 matched samples, GEOMeta disease-status assignments were concordant with DiSignAtlas case/control labels for 96.23%. Among 15,179 samples with tissue annotations available in both resources, 96.95% showed concordant annotations. Next, we compared CP annotations with PharmGEO^29^, a manually curated resource focused on drug-induced transcriptomic profiles. Among 10,359 overlapping samples, compound names were concordant for 96.34%, PharmGEO treated/control assignments agreed with GEOMeta perturbation type for 95.48% and with sample-level perturbation status for 96.44%. Detailed match-category breakdowns, status comparisons, and discordant cases for the DiSignAtlas and PharmGEO analyses are provided in Supplementary Results 1.1 and 1.2, respectively.

Because sex and age lack manually curated reference datasets, we evaluated consistency against manually reviewed evidence from the original GEO metadata. Cases where both extraction methods yielded missing data were analyzed separately, which accounted for 77.78% of samples for sex and 83.55% for age. Among samples with non-missing annotations from both approaches, agreement was 99.96% for sex and 95.51% for age. GEOMeta identified sex and age information in more samples than conservative text-based extraction, reflecting its ability to integrate evidence distributed across multiple metadata fields and free-text descriptions.

To evaluate whether transcriptome-based models could contribute to metadata auditing, we selected the sex-prediction tasks and treated discrepancies between model predictions and original metadata labels as auditing signals. Among 3,551 samples in the 2024 holdout, 173 were label-prediction discordant, including 57 female-labeled samples predicted as male and 116 male-labeled samples predicted as female. In both groups, sex-marker gene expressions shifted toward the predicted sex (Fig. 4C and D). Raw-read auditing of ten selected female-labeled discordant samples further identified five with male-associated transcriptional signals, four with indeterminate profiles, and one with a female-associated profile (Supplementary Data S1, see the Sex Prediction Section for details). Thus, model disagreement successfully prioritized biologically informative and technically problematic samples, but did not uniformly indicate metadata error, supporting its use as a diagnostic signal for targeted review rather than as ground truth.

### Benchmarking in Transcriptome Representation Learning

By linking harmonized sample-level annotations to bulk RNA-seq expression profiles, GEOMeta enables reproducible label-prediction tasks under consistent evaluation settings. We used samples deposited before 2024 for model training, while reserving samples deposited in or after 2024 as a prospective temporal holdout (2024-holdout). In a separate study-level holdout evaluation, we randomly selected entire studies from the pre-2024 dataset as the test set (study-holdout), ensuring that all samples from a given study were assigned exclusively to either the training or test set. We considered four representative tasks: prediction of sex, tissue, disease and age group. Although related tasks have been examined previously^45–48^, most studies have relied on relatively homogeneous cancer and normal-tissue datasets such as TCGA and GTEx. Our goal was not to introduce new modeling methods, but to provide a large, diverse and carefully curated benchmark for evaluating transcriptomic representation-learning approaches. Notably, unlike previous efforts, the extensive diversity of diseases represented within our blood samples enabled us to evaluate disease classification models using blood transcriptomes alone. This provides a unique benchmark for developing minimally invasive, blood-based diagnostic models.

To establish baseline results, we compared a diverse set of modeling strategies that span both bulk RNA-seq-specific approaches and foundation models originally trained on single-cell transcriptomic data. Specifically, we included BulkFormer^9^, a transformer model recently pretrained on bulk transcriptomes, together with single-cell-derived representation models including scGPT^10^, CellPLM^13^, and scVI^49^. For the foundation models, we evaluated two transfer-learning settings where applicable: frozen-encoder with MLP probing, in which the pretrained encoder is used as a fixed feature extractor and only a downstream MLP classifier is trained, and end-to-end fine-tuning, in which the pretrained model and classification head are jointly optimized for each prediction task. This design allows GEOMeta to benchmark not only model families, but also different adaptation strategies for applying pretrained models to heterogeneous bulk RNA-seq data.

Although scGPT and CellPLM were originally developed using single-cell transcriptomic data, they may still provide useful representations for bulk RNA-seq. Single-cell foundation models can learn gene-gene co-expression structure, pathway-level relationships, and regulatory patterns from large numbers of individual cells at the scale of 100 million cells ^10,50^. Many of these relationships are not restricted to one specific cell type, but reflect broader transcriptional programs shared across cellular contexts^51^. A bulk RNA-seq profile can be viewed, for representation-learning purposes, as an aggregate pseudo-cell: a single gene-expression vector formed by pooling signals across many cells in a tissue or sample. Although cell-type-specific signals may be diluted in bulk data, conserved gene-gene relationships and shared biological programs can remain detectable. This provides a rationale for comparing single-cell pretrained encoders with bulk-specific models on GEOMeta.

Prediction performance differed markedly across tasks (Fig. 3A). Sex prediction achieved the highest accuracy, consistent with the strong signals contributed by sex-linked genes. Tissue prediction also performed well, reflecting robust tissue-specific expression programs. In contrast, disease prediction was more difficult because disease labels are biologically heterogeneous and confounded by tissue of origin, cellular composition and study design. For example, blood samples from patients with lung cancer may not capture the transcriptomic state of the primary tumor and T cells isolated from brain tissues may not fully capture the pathological processes underlying Alzheimer’s disease. Notably, the trained models more frequently failed to distinguish disease samples from normal controls rather than from another disease. Age-group prediction showed the lowest performance, consistent with the subtle, continuous and context-dependent nature of age-associated transcriptional changes in heterogeneous bulk RNA-seq data sets.

**Figure 3:**
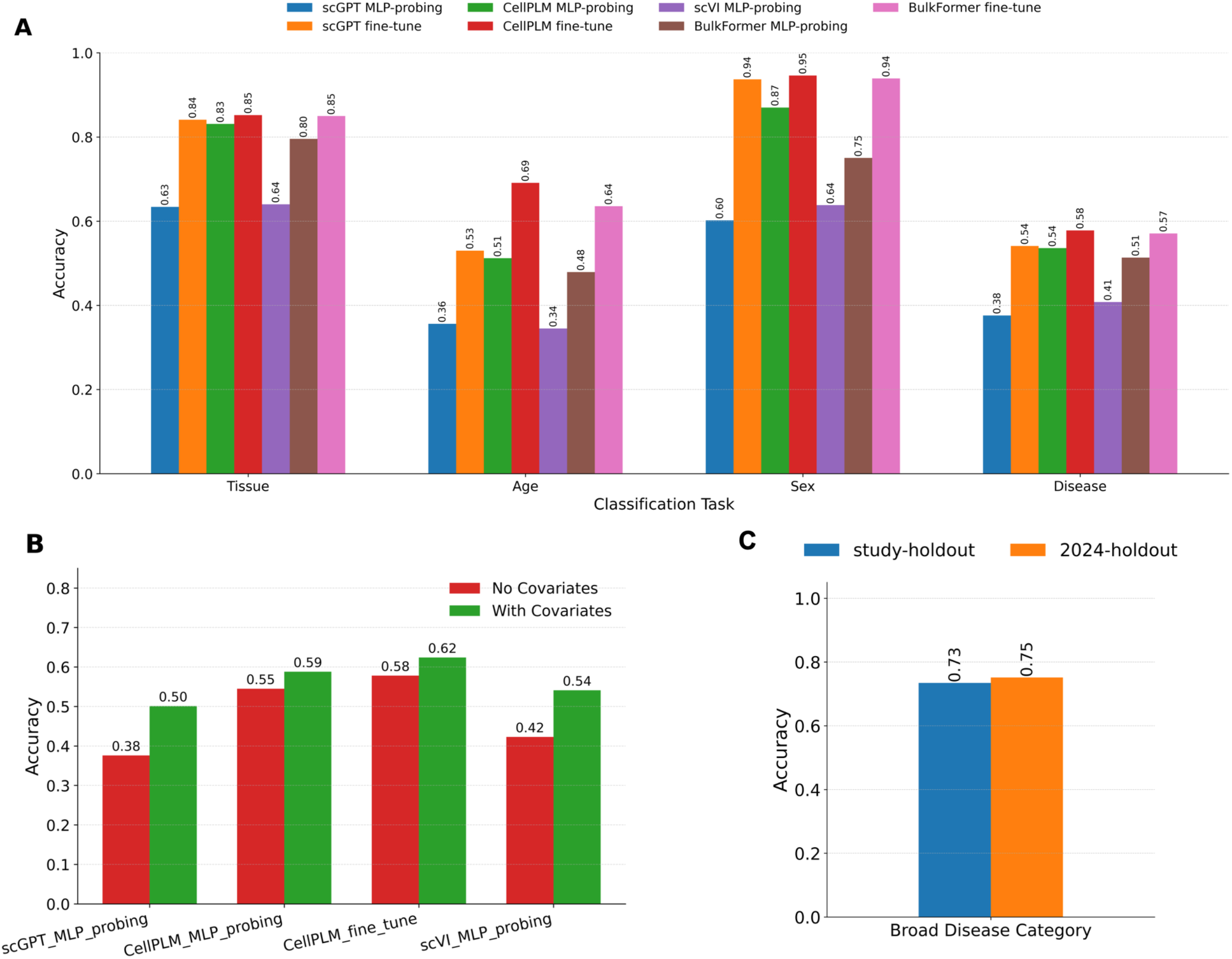
Benchmarking in transcriptome representation learning in meta feature prediction. A: Label prediction results for the 2024 holdout dataset. Accuracy achieved by seven methods scGPT (MLP-probing), scGPT (Fine-Tune), CellPLM (MLP-probing), CellPLM (Fine-Tune), scVI (MLP-probing), Bulkformer (MLP-probing), Bulkformer (Fine-Tune) across four prediction tasks (Sex, Organ, Disease, and Age). **B**: Disease prediction results for the 2024 holdout datasets. Accuracy achieved by four methods scGPT (MLP-probing), CellPLM (MLP-probing), CellPLM (Fine-Tune), scVI (MLP-probing) with and without additional covariate information. **C**: Broad disease group classification accuracy for CellPLM (Fine-Tune) across three test sets.

Across tasks, fine-tuned models generally outperformed zero-shot encoder representations evaluated with MLP probes (Fig. 3A), consistent with previous reports ^52^ and indicating that task-specific adaptation is important when transferring single-cell foundation models to bulk RNA-seq data. Fine-tuned CellPLM achieved the strongest overall performance, suggesting that representations learned from large-scale single-cell datasets retain biologically meaningful information in bulk transcriptomes. Nevertheless, the lower performance for disease and age prediction highlights persistent challenges arising from tissue heterogeneity, batch effects, study-specific designs and label granularity.

To improve disease prediction, we incorporated sex, tissue and experimental setting as covariates. Each covariate was embedded and combined with the representation generated by the pretrained foundation model, and the resulting representation was passed to an MLP classifier. Incorporating these contextual features improved disease-prediction accuracy (Fig. 3B and Supplementary Fig. S1), consistent with the tissue specificity and sex dependence of many diseases.

As the disease classification achieved only modest accuracy across methods (50%-64%; Fig. 3A and Supplementary Fig. S1), we evaluated whether the models could resolve broader clinical categories, including oncology, immune, neurological, digestive and infectious diseases. This analysis was restricted to blood-derived samples and retained normal controls as the largest class, yielding 11 categories in the 2024-holdout and 12 in the study-holdout. Performance improved markedly under this broader taxonomy (Fig. 3C). Fine-tuned CellPLM achieved accuracies of 75% and 73% in the 2024 and study holdouts, respectively, Thus, blood transcriptomes contain robust signals that distinguish broad disease classes from one another and from healthy controls. The pronounced gap between broad disease category and disease classification suggests that the main remaining challenge is resolving within-category heterogeneity, such as distinguishing individual cancer and infection types, in the presence of tissue-of-origin differences and study-specific technical effects.

Although age-group classification accuracy was modest, the confusion matrix showed that errors were concentrated near the diagonal rather than distributed randomly (Supplementary Fig. S2). Most misclassifications occurred between adjacent age groups, consistent with the continuous nature of aging; for example, 273/758 (36.0%) Adult samples were classified as Late Adult/Elderly, but only 33/758 (4.3%) were classified as Youth. This pattern suggests that the model captured the ordinal structure of age-related transcriptomic variation even when it failed to assign the exact category. We therefore additionally evaluated performance using Spearman correlation, encoding Youth, Adult and Late Adult/Elderly as 0, 1 and 2, respectively (Supplementary Fig. S1) and observed a significant positive correlation between the predicted and true age groups.

### Sex Prediction Reveals Biological Determinants and Flags Discordant Samples

Sex prediction was the most accurate task and has a well-characterized transcriptomic basis, enabling us to examine both model interpretability and its potential for metadata quality control. We first compared the fine-tuned models with a two-layer multilayer perceptron trained using only 35 established sex-associated genes, comprising 34 Y-chromosome genes and female-specific marker *XIST*. This marker-based model achieved accuracies of 95% on the 2024 holdout compared with 95% for fine-tuned CellPLM (Fig. 4A). Thus, biological sex in bulk RNA-seq is largely captured by a compact and well-established transcriptional signature.

**Figure 4:**
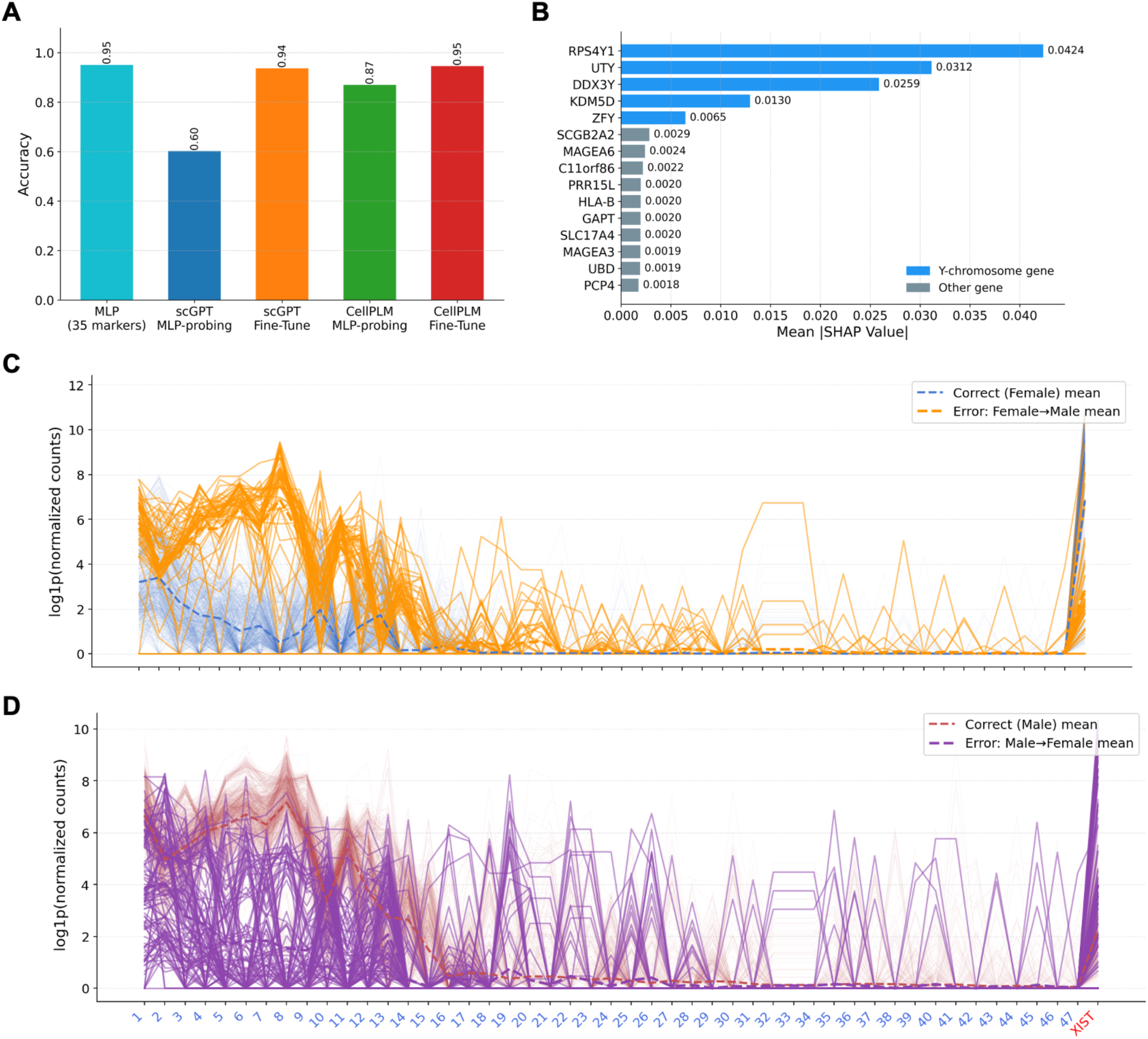
Results on sex prediction. **A:** Sex prediction accuracy of a marker-based MLP baseline versus single-cell foundation models. Grouped bar chart comparing sex classification accuracy on the 2024 holdout datasets across six approaches: a simple two-layer MLP trained on 35 known sex-associated marker genes (34 Y-chromosome genes and XIST), and four foundation model methods (scGPT MLP-probing, scGPT Fine-Tune, CellPLM MLP-probing, CellPLM Fine-Tune). **B:** Top 15 genes ranked by mean absolute SHAP value for male prediction using CellPLM fine-tuned; Y-chromosome genes are shown in blue and other genes in gray. Each polyline represents the expression of one sample across 48 sex-linked marker genes. **C:** Female-labeled samples: concordant (*n* = 1,964, blue) vs. misclassified as male (*n* = 57, orange, bold). **D:** Male-labeled samples: concordant (*n* = 1,414, red) vs. misclassified as female (*n* = 116, purple, bold). Dashed lines indicate group means. Gene identities for each number are provided in Supplementary Fig. S3. *XIST* (red, rightmost) is the sole X-chromosome marker gene included.

Consistent with this result, SHAP analysis^55^ of fine-tuned CellPLM identified canonical Y-chromosome genes as the dominant predictive features (Fig. 4B). *RPS4Y1*, *UTY*, *DDX3Y*, *KDM5D,* and *ZFY* had the highest mean absolute SHAP values, with contributions approximately an order of magnitude greater than those of other genes. Although these markers were not explicitly supplied as prior knowledge, the model recovered the expected molecular determinants of biological sex, supporting the biological fidelity of both the learned representation and the GEOMeta annotations.

We next examined sex-marker expression in correctly and incorrectly classified samples (Fig. 4C, Fig.4D). Misclassifications were concentrated among samples with atypical marker profiles rather than distributed randomly. Female-labeled samples predicted as male showed elevated Y-chromosome gene expression resembling the canonical male profile, whereas male-labeled samples predicted as female showed reduced Y-linked expression together with elevated *XIST*. In both groups, the metadata–prediction discordant samples shifted toward the expression profile of the predicted sex, consistent with the SHAP-derived feature importance.

To determine whether the Y-chromosome signal in the discordant female-labeled samples predicted as male could be explained by ambiguous read mapping, we reanalyzed raw RNA-seq data from 16 SRA runs representing 14 unique GEO samples. The audit cohort comprised four concordant control samples (2 female, 2 male), and 10 discordant female-label samples (two discordant samples were represented by two SRA runs each). The concordant reference samples showed the expected reciprocal expression patterns: concordant female-labeled samples exhibited abundant *XIST* expression and negligible chr Y-linked transcription, whereas concordant male-labeled samples exhibited robust expression of multiple canonical chr Y-linked genes and no detectable *XIST* (Supplementary Data S1). These observations established baseline transcriptional profiles and supported the validity of the alignment and quantification procedures.

Among the 10 unique discordant samples, five were classified as exhibiting male-associated transcriptional signal, four were indeterminate, and one retained a female-associated profile. In the three strongest male-associated cases (GSM4061943, GSM5656873, and GSM5683376), multiple canonical Y-linked genes, including *RPS4Y1*, *DDX3Y*, *KDM5D*, and *UTY*, were coordinately expressed, while *XIST* was absent or nearly absent. Moreover, 91.9-98.2% of chr Y-aligned reads in these samples were uniquely mapped. Furthermore, the ratio of uniquely mapped chromosome Y reads to total genome-wide uniquely mapped reads in these samples was consistent with biological males, representing an order-of-magnitude increase over the background noise observed in female controls. This strong concordance between expression profiles, high-confidence alignments, and genome-wide read proportions firmly argues against homologous multimapping as the primary driver of the observed Y-linked signal.

The four indeterminate samples underscore the necessity of integrating sequencing-quality metrics with model-guided audits. For example, GSM5357913, GSM8023018, and GSM8022984 contained fewer than six million total mapped reads per run and exhibited low genome-wide unique mapping rates, which limits the reliable detection of both *XIST* and Y-linked markers, rendering them ambiguous. Meanwhile, GSM8263754 displayed substantial *XIST* expression alongside Y-linked transcription, indicating a mixed or biologically complex sex-chromosome profile. Conversely, both SRA runs for GSM8398175 demonstrated reproducible *XIST* expression, weak Y-marker expression, and extremely low ratios of uniquely mapped chromosome Y reads to genome-wide reads, ultimately supporting a female-associated transcriptional profile despite the model’s initial prediction. Thus, not every prediction-annotation disagreement stems from a metadata error. Model predictions serve as powerful diagnostic flags to prioritize samples for targeted, sequence-level review. Together, these results show that transcriptome-based prediction not only recovers established biology but also provides a sensitive, model-guided approach for identifying potentially erroneous or biologically atypical samples in large transcriptomic collections.

### Deployment of the annotation pipeline to prospective datasets

Because GEOMeta was primarily developed using studies available in ARCHS4 v2.5 (January 13, 2025), we next evaluated its performance on newly submitted GEO datasets. We first analyzed a selected sample of 96 GEO Series comprising 7,319 samples submitted in May 2026. Using GPT-5, GEOMeta completed annotation in approximately 15 hours at an estimated API cost of US$53. A domain expert manually reviewed annotations for 1,000 samples. Disease, tissue and age annotations were confirmed for all reviewed samples, whereas sex annotations were confirmed for 99.7% of samples. All reviewed chemical-perturbation annotations were also confirmed (Supplementary Table S3). Although these results demonstrated that prospective automated curation was feasible, the computational cost remained substantial (approximately US$0.50 per one GEO Series), motivating a systematic evaluation of newer and more economical LLMs.

We therefore benchmarked 22 frontier LLMs from 11 providers using a benchmark of 118 samples from five GEO Series submitted by our laboratory. Models were evaluated under identical inputs for run completion, final-output completeness, concordance with a cross-model consensus, runtime and API cost. After normalization, estimated runtime for a typical six-sample study ranged from 0.18 to 2.38 minutes, while estimated API cost ranged from US$0.0013 to US$0.0666 (Fig. 5 and Supplementary Data S2).

**Figure 5.**
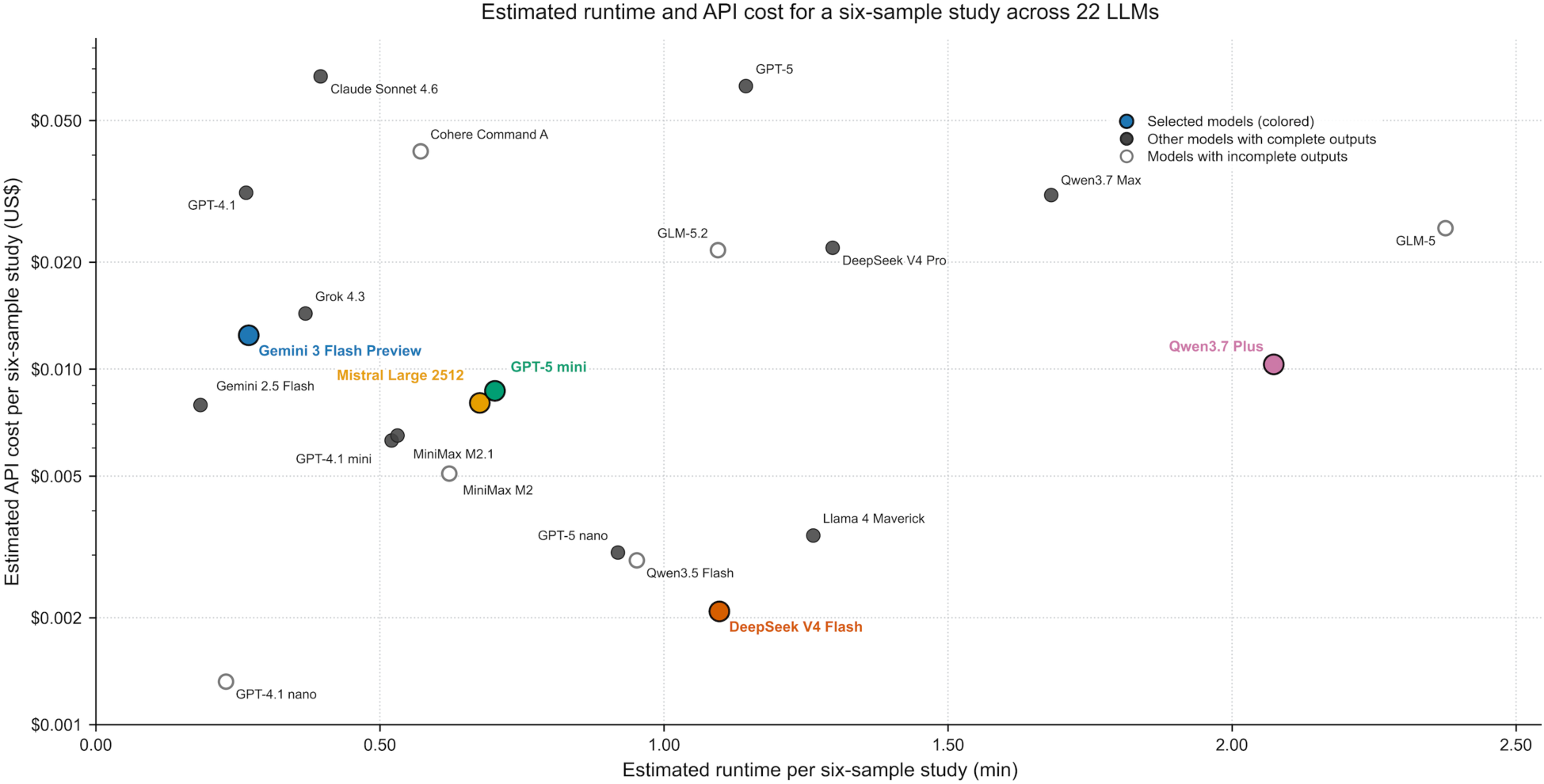
Internal multi-LLM benchmark of runtime and API cost. Estimated runtime and API cost for a typical six-sample study across 22 LLMs using the internal 118-sample benchmark. Colored points indicate the five models selected for prospective evaluation; filled gray points indicate other models with complete outputs, and open points indicate models with incomplete outputs.

As expected, frontier reasoning models such as GPT-5 and Claude 4.6 incurred substantially higher costs than lightweight open-source models such as Qwen Flash and DeepSeek Flash, with approximately 21- to 32-fold differences in API cost. Gemini 2.5 Flash achieved the fastest estimated runtime, whereas GPT-4.1 nano had the lowest estimated API cost but did not produce complete outputs for all benchmark samples. In contrast, GLM-5 and GLM-5.2 generated numerous missing annotations despite their strong coding performance, suggesting reduced effectiveness for scientific data inference. Based on annotation completeness, consensus agreement, runtime and API cost, we selected five models including Gemini 3 Flash Preview, Mistral Large 2512, GPT-5 mini, DeepSeek V4 Flash and Qwen3.7 Plus for the prospective evaluation. All five generated complete outputs for both the 118-sample benchmark and the 18-sample CP subset. Concordance with the benchmark consensus ranged from 92.8% to 97.9%, whereas PubChem-mapped CP identities achieved 100% agreement in the CP benchmark.

Because few May 2026 studies had processed expression profiles available in the current ARCHS4 release, we assembled a separate expression-linked cohort from December 2025 to January 2026 for the multi-model prospective evaluation. The cohort comprised 225 GSEs, including 5,374 samples with matched expression profiles and 6,733 associated GSM records retrieved for annotation. Among the five selected models, the relative runtime ordering and broad API cost pattern were consistent with those observed in the internal benchmark. Estimated runtime for a typical six-sample study ranged from 0.43 minutes for Gemini 3 Flash Preview to 1.16 minutes for Qwen3.7 Plus, while estimated API cost ranged from US$0.0025 for DeepSeek V4 Flash to US$0.0146 for Gemini 3 Flash Preview (Fig. 6A and Supplementary Table S4). The cost pattern was also broadly consistent: DeepSeek V4 Flash remained the least expensive model, Gemini 3 Flash Preview remained the most expensive, and the other three models formed an intermediate-cost group, although their exact ordering differed. Across the five models, normalized API cost estimates were 9.6-46.4% higher in the prospective cohort, with a mean increase of 22.8% and absolute differences below US$0.004 per typical six-sample study. These results supported extrapolation of the broad runtime and cost patterns observed in the internal benchmark to the larger prospective cohort.

**Figure 6.**
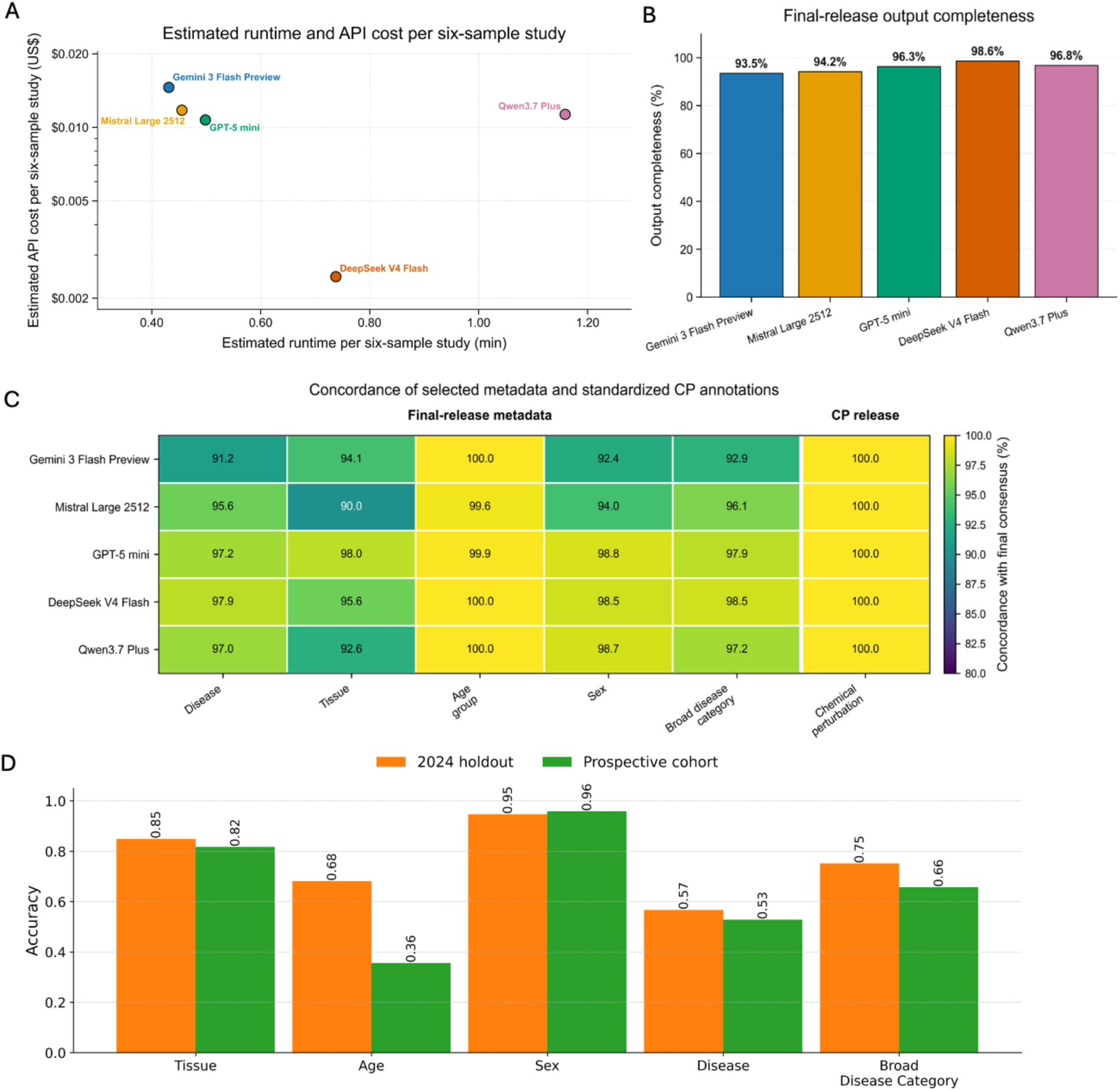
Prospective multi-LLM evaluation and downstream label-prediction performance. **A.** Estimated runtime and API cost for a typical six-sample study for the five selected models applied to 6,733 GSM records from 225 GEO Series. Per-sample runtime and API cost were calculated from the total cohort values and multiplied by six to obtain estimates for a typical six-sample study. **B.** Final-release output completeness for the five selected models in the prospective multi-LLM cohort. **C.** Concordance of selected metadata fields with the final cross-model consensus across the five models, together with concordance of PubChem-mapped chemical identity in the CP-focused release. **D.** Accuracy of fine-tuned CellPLM across five classification tasks, tissue, age, sex, disease and broad disease category on the 2024 temporal holdout (orange) and the December 2025-January 2026 cohort (green, “prospective cohort”).

Final-release output completeness ranged from 93.5% for Gemini 3 Flash Preview to 98.6% for DeepSeek V4 Flash (Fig. 6B). The final consensus release contained 5,945 samples. Across the five models, concordance with the final consensus ranged from 91.2%-97.9% for disease, 90.0%-98.0% for tissue, 92.4%-98.8% for sex, 92.9%-98.5% for broad disease category, and was highest for age group, at 99.6-100.0% (Fig. 6C). The final CP-focused release contained 842 samples across 41 GSEs, and PubChem-mapped chemical identity showed 100% concordance among available model outputs in the retained cohort. After restriction to human bulk RNA-seq samples with available expression profiles, 4,647 samples remained for evaluation of transcriptomic representation-learning methods.

To assess whether the representation-learning results generalized beyond the 2024 holdout, we applied the best-performing model, fine-tuned CellPLM, to the prospectively annotated December 2025-January 2026 cohort (“prospective cohort”) across five classification tasks (Fig. 6D). Performance was broadly preserved between the two evaluation sets. Sex prediction remained the most accurate and was essentially unchanged (95% versus 96%), and tissue prediction stayed high with only a marginal decrease (85% versus 82%). Fine-grained disease classification showed a similarly small reduction (57% versus 53%), and broad disease category classification again outperformed fine-grained disease on both sets (75% versus 66%), consistent with our earlier observation that broader clinical categories are more readily resolved from bulk transcriptomes. Age-group prediction was the notable exception, falling from 68% on the 2024 holdout to 36% on the prospective cohort. This pronounced decline reinforces that age is the least robust and most context-dependent of the evaluated tasks, in line with the subtle and continuous nature of age-associated transcriptional variation and its sensitivity to differences in cohort composition across release periods.

Together, these results demonstrate that recent open-source models, particularly DeepSeek V4 Flash, can achieve annotation quality comparable to that of leading frontier models while reducing inference costs by more than an order of magnitude, making large-scale automated curation of public transcriptomic repositories economically practical.

## Discussion

GEO remains an invaluable resource for transcriptomic research, yet its large-scale reuse continues to be constrained by heterogeneous and inconsistently structured metadata. In this work, we developed GEOMeta, which makes two contributions: 1) a quality-controlled, sample-level metadata resource comprising nearly 600,000 human bulk RNA-seq samples and 2) a reusable multi-stage workflow for annotating, standardizing, and validating GEO metadata. To our knowledge, this represents one of the largest quality-controlled, sample-level metadata resources for human bulk RNA-seq studies. Importantly, GEOMeta performs annotation at the individual sample level rather than assigning labels only at the study level, enabling sample-wise dataset reuse within and across GEO series. More broadly, this work positions metadata standardization not simply as a preprocessing task, but as a reusable infrastructure layer for scalable transcriptomic analysis and reuse.

A central observation is that metadata annotation quality is largely determined during the extraction stage. While constrained fields (e.g., sex, age) were robust to different prompt designs, context-dependent domains (e.g., disease, tissue, perturbation) were highly sensitive. Generic extraction increased missingness for tissue and perturbation and produced broader or more fragmented labels. For disease, lower missingness was accompanied by reduced biological specificity. These extraction-stage differences frequently persisted through downstream standardization and ontology mapping.

More broadly, large-scale GEO metadata curation extends beyond simple entity extraction. The central challenge is not simply identifying biomedical terms, but determining which attributes apply to each biospecimen from information distributed across study- and sample-level records. Once these sample-specific attributes are identified, they must be expressed consistently, converted into secondary labels under explicit rules, and linked to appropriate ontology terms. GEOMeta thus separates contextual extraction, normalization, controlled label derivation, and ontology alignment into modular and independently reviewable stages. This separation makes errors traceable and allows individual components to be evaluated or updated independently. It is important because information missed or misassigned during extraction cannot be reliably recovered through downstream normalization and ontology mapping. Also, by combining LLM-based contextual interpretation with rule-guided normalization and curated ontology mapping, GEOMeta provides a hybrid annotation framework that improves consistency while preserving biologically meaningful interpretation across heterogeneous metadata sources. Together, these design choices contribute substantially to the robustness, interpretability, and reproducibility of large-scale metadata curation compared with treating the entire annotation task as a single unconstrained extraction step.

Beyond metadata curation, the downstream benchmarking results illustrate the biological and methodological limits of current transcriptome representation models. Sex prediction achieved high performance, whereas disease and age prediction remained substantially more difficult. This pattern is biologically plausible. These results indicate that prediction difficulty partly reflects limitations in biological signal and metadata structure rather than model architecture alone. Disease categories are often broad, clinically heterogeneous, and partly confounded by tissue of origin, experimental setting, and sample composition. Age-related transcriptomic effects are also relatively subtle in bulk data and can be obscured by cell-type mixture and study design differences. The improvement observed when sex, tissue, and experimental setting were incorporated as covariates further supports the view that transcriptome-based prediction is strongly context dependent. These results suggest that future transcriptome models may benefit from explicitly using metadata structure rather than treating prediction targets as independent flat labels. Hierarchical disease labels, ontology-aware learning, and multi-task frameworks that jointly model disease, tissue, age, sex, and experimental context may better reflect the structure of the data.

The prospective comparison also provided practical guidance for model selection in routine deployment. Runtime and API costs scaled consistently between the small and median-size cohorts, providing a practical basis for approximate budget and runtime estimation for future annotation projects. Although frontier reasoning models may offer advantages for complex reasoning tasks, our results indicate that their higher inference costs are not necessarily justified for large-scale metadata annotation. With carefully designed agents and task-specific prompts, lower-cost Flash models achieved completeness and consensus concordance comparable to higher-cost reasoning models at a small fraction of the cost. These findings highlight the potential of efficient open-source LLMs for scalable scientific curation, particularly in settings involving sensitive or privacy-restricted data where self-hosted deployment is desirable.

This study has several limitations. First, the absence of comprehensive gold-standard annotations remains a fundamental challenge for large-scale GEO metadata curation. GEO does not provide validated reference annotations for most metadata domains, and apparent discrepancies may reflect ambiguity, incompleteness, or inconsistency in the original submissions rather than annotation failure alone. More broadly, the quality of any curation framework is ultimately constrained by the quality of the underlying source metadata. GEO studies vary substantially in reporting completeness, terminology usage, and metadata granularity. Although GEOMeta integrates information across multiple metadata sources and incorporates expert-reviewed quality control procedures, missing or incorrect information in the original records cannot always be resolved automatically. Missing annotations may reflect either unreported information or information that was insufficiently specific for confident resolution.

Second, annotation confidence is not uniform across metadata domains. Certain fields, such as sex and age group, are often reported explicitly and can frequently be derived from structured metadata. In contrast, disease, perturbation, and experimental-setting annotations commonly require integration of evidence distributed across study descriptions, sample annotations, and supporting publications. As a result, residual ambiguity may remain for studies with incomplete or inconsistently reported metadata, and not every curated sample is equally suitable for every downstream application. Effective sample size therefore depends on field-specific completeness, annotation confidence, and the requirements of a particular analytical task.

Third, the multi-LLM consensus used in the model comparison represents agreement among models rather than an independent accuracy standard. Models may share correlated errors, and high consensus agreement does not guarantee biological correctness. Consensus-based results should therefore be interpreted primarily as measures of output stability and cross-model consistency. Independent expert review remains necessary for evaluating difficult or high-impact annotation domains.

Fourth, GEOMeta primarily focuses on sample-level biological annotation and does not explicitly represent higher-order study design structures. Many GEO studies contain experimental designs that extend beyond simple disease-versus-control or perturbation-versus-control comparisons, including paired samples, longitudinal sampling, matched cohorts, multifactorial interventions, and cell-type-specific profiling strategies. Although some of this information may be present within study descriptions and sample metadata, it is often distributed across multiple metadata fields and is not consistently reported in a structured format. Consequently, GEOMeta annotations should not be interpreted as a complete representation of study design, and users may need to review study-level metadata when constructing disease signatures, perturbation signatures, or other specialized downstream analyses.

Finally, GEOMeta is currently optimized for a focused set of metadata fields in human bulk RNA-seq studies. Future extensions could encompass additional attributes, including strain, treatment response and survival, as well as other modalities such as CITE-seq and ChIP-seq. Each new field and assay type will require tailored prompts and rigorous validation to maintain annotation quality. Nevertheless, our results demonstrate that the language-processing and reasoning capabilities of current LLMs are already sufficient to support scalable, automated scientific curation.

## Online Methods

### Manual Curation and Preliminary Assessment

Before developing the large-scale annotation pipeline, we performed several rounds of manual curation to understand how GEO metadata are reported in practice. During the COVID-19 pandemic, four medical students manually annotated age, tissue, and sex for 20,000 human samples in a study of sex-associated differences in COVID-19. In a later effort, three high-school interns received task-specific training and collectively curated metadata for an additional 13,000 samples over two months. They first screened GEO Series to identify eligible drug-induced bulk gene-expression studies and then annotated approximately 10-12 metadata fields, including age, sex, tissue and perturbation-related attributes. The resulting annotations were reviewed and standardized by a pharmacology graduate student within 15 days. These efforts were useful for defining annotation rules, but also demonstrated the limitations of manual curation. Metadata were often incomplete, inconsistently formatted, or distributed across multiple GEO fields. Even after training, curators differed in how they interpreted ambiguous sample descriptions, particularly for tissue source, disease status, and perturbation conditions. Manual review was therefore helpful for developing annotation guidelines but was not scalable to hundreds of thousands of samples.

We also evaluated MetaMap^54^ as a text-mining tool for GEO metadata. While it could identify some biomedical terms, its reliance on direct concept matching made it difficult to resolve cases requiring study context, such as distinguishing tissue source from disease state or identifying whether a treatment term referred to a perturbation or a control condition. These observations motivated the development of a structured annotation pipeline that combines automated extraction, field-specific normalization, and targeted review.

### Problem Characterization

Large-scale GEO metadata annotation presents challenges beyond conventional text extraction because biologically relevant information is frequently fragmented across metadata fields, inconsistently represented across studies, and dependent on experimental context. In addition, different metadata domains (e.g., disease, tissue, age, sex, perturbation) exhibit distinct sources of ambiguity and require different forms of interpretation and normalization. Based on manual curation and preliminary tool evaluation, we identified several recurring sources of difficulty in GEO metadata annotation. First, biologically relevant information is rarely contained within a single metadata field. Study-level descriptions (GSE) often summarize overall experimental design, whereas sample-level metadata (GSM) contain sample-specific attributes. Accurate annotation therefore requires integrating information across both levels rather than relying on isolated fields.

Second, metadata interpretation depends strongly on study context. The same term may represent different biological meanings depending on sample origin, disease setting, or experimental design. For example, terms such as “normal” may refer to healthy tissue, adjacent normal tissue from diseased donors, or untreated baseline conditions. Similarly, tissue descriptors, mutation labels, and treatment annotations may be incorrectly interpreted when evaluated without surrounding study context. Certain metadata attributes also require implicit inference rather than direct extraction. For example, donor sex, disease status, or perturbation conditions may not be explicitly stated within structured metadata fields and instead require interpretation of study descriptions, sample naming conventions, mutation backgrounds, or linked publications. As a result, accurate annotation often depends on integrating multiple indirect sources of evidence rather than relying solely on explicit metadata labels.

Different metadata domains also exhibit distinct forms of ambiguity and heterogeneous representation. Age fields may describe donor age, developmental stage, or experimental timing, whereas tissue annotations may combine anatomical structures, cell types, developmental regions, and culture systems within the same field. Disease annotations are similarly heterogeneous and frequently distributed across study descriptions, sample annotations, and linked publications rather than represented as standardized sample-level labels.

Age annotation was a clear example of this challenge. GEO age fields may describe human donor age, fetal developmental age, or non-donor timing such as culture duration, differentiation time, treatment time, passage number, or animal host age. Correct interpretation therefore depends on both units and study context. Numeric values without sufficient context can also be misleading. For example, the metadata for GSM5942992 reported “age: 50.” If interpreted in isolation, this could incorrectly suggest 50 years of age. However, the associated GSE metadata indicated that blood was collected from healthy infants at week 6 and week 7 of life, supporting interpretation as 50 days of age. In other cases, age information was distributed across multiple metadata fields rather than provided directly. Examples included gestational descriptions embedded in source names, ages reported in sample titles, or biospecimen ages derived from date fields. Conversely, values describing experimental timing, passage number, or non-human host age could not be interpreted as human biological age. Representative raw-to-final annotation examples are summarized in Supplementary Table S5.

Tissue annotation presented similar context-dependent challenges. GEO tissue fields frequently contain mixtures of anatomical organs, tissue subregions, developmental structures, cell types, disease sites, culture systems, and xenograft implantation sites. Therefore, reliable annotation required distinguishing the reported site from the underlying tissue of origin and determining when a specific descriptor could be mapped to a broader controlled category. Representative examples are provided in Supplementary Table S6.

Disease annotation introduced additional complexity because disease information in GEO is rarely provided as a single standardized sample-level field. Relevant evidence may instead appear across GSM characteristics, sample titles, source descriptions, GSE titles, study summaries, overall designs, and linked publications. GEO studies may also contain multiple biological groups within the same series, including diseased samples, healthy controls, adjacent normal tissues, disease subtypes, or progression states. Terms such as “control,” “normal adjacent tissue,” or “recurrent tumor” may represent biologically distinct disease contexts despite superficially similar wording. Disease labels may also vary in specificity, ranging from broad categories such as “brain tumor” to specific histological or molecular subtypes. As a result, disease labels cannot be assigned reliably using study-level descriptions alone and require context-aware interpretation of both study-level and sample-level metadata.

Additional challenges arise from studies with complex experimental designs that are not easily represented by a small set of standardized annotation fields. Examples include paired case-control studies, longitudinal sampling designs, isolated cell-population profiling, ex vivo culture systems, and multifactorial perturbation experiments. In these settings, biologically meaningful interpretation often depends on relationships among samples rather than attributes of individual samples alone. Furthermore, procedures used for sample collection, cell isolation, or culture may resemble perturbations at the metadata level despite not representing true biological interventions. These scenarios illustrate the limitations of direct keyword-based annotation and motivated the development of a staged framework integrating study-level context, sample-level metadata, standardization procedures, and expert-reviewed quality control.

Together, these observations indicate that GEO metadata annotation cannot be reliably addressed through isolated field parsing or direct string matching alone. Effective annotation requires context-aware integration of study-level and sample-level metadata, semantic normalization across heterogeneous reporting styles, and structured ontology-aware interpretation. These challenges motivated the development of a staged annotation framework separating extraction, refinement, controlled derivation, standardization, and ontology mapping into independently reviewable steps.

### Task Overview

The objective of this work was to transform unstructured GEO metadata into structured, standardized, and machine-readable annotations at the sample level. For each GSM-GSE pair, GEOMeta generated a structured annotation record integrating both study-level and sample-level metadata. In total, the annotation schema comprised 27 metadata fields reflecting the range of descriptors commonly reported in GEO submissions (Supplementary Table S7).

In this study, we focused on a subset of fields broadly used in downstream modeling and comparative transcriptomic analyses: RNA library type, RNA source, experimental setting, study-level perturbation, sample-level perturbation, disease, tissue, age group, and sex. The chemical-perturbation analyses additionally included perturbation identity, perturbation type, dosage and duration. These fields capture major biological and experimental sources of variation in bulk RNA-seq data and form the basis for the structured labels used in subsequent analyses. Figure 1 summarizes the GEOMeta resource, the multi-stage annotation framework, and its prospective application to newly released GEO studies. Specifically, Figure 1A presents the released sample-level transcriptomics metadata resource and its intended downstream applications, Figure 1B the annotation, quality-control, standardization and mapping workflow, and Figure 1C illustrates the prospective application to GEO studies submitted in December 2025 and January 2026.

### Data Retrieval and Preprocessing

To curate a high-quality set of human bulk RNA-seq samples, we applied a multi-step filtering pipeline. We first extracted human samples from the ARCHS4 reference file (*human_gene_v2.5.h5*), which contained 888,821 RNA-seq samples across multiple experimental types and platforms, including both bulk and single-cell assays. Each sample is then assigned a probability of being single-cell; samples with a probability below 0.1 were retained as bulk, yielding 490,401 initial samples. Because a single GSM accession can map to multiple GSE identifiers, we expanded GSM-GSE relationships and obtained 592,367 GSM-GSE pairs. SuperSeries were removed due to lacking detailed study design, while their corresponding SubSeries were retained. After excluding 101,788 SuperSeries samples, the final ARCHS4-derived set contained 490,579 valid GSM-GSE pairs, slightly greater than the original 490,401 GSMs, reflecting cases where a GSM was associated with more than one valid SubSeries. Metadata for these candidates was programmatically retrieved from GEO accession pages in XML format using Python, and both series-level (GSE) and sample-level (GSM) information were parsed into structured records. Additional exclusions included private accessions not yet released on GEO, accessions marked as deleted by GEO staff, and samples that consistently failed XML retrieval across up to 6 attempts.

To increase coverage beyond ARCHS4-derived samples, we additionally queried GEO directly using the search string *“homo sapiens [Organism] AND expression profiling by high throughput sequencing [DataSet Type]*”. This yielded an additional 228,250 GSM-GSE pairs. Metadata for these samples was processed with the filtering pipeline to exclude non-bulk modalities such as single-cell, ChIP-seq, ATAC-seq, methylation, and ribosome profiling. Merging ARCHS4-derived and directly retrieved samples produced a combined set of 718,829 GSM-GSE pairs for annotation.

Following metadata annotation, we excluded records inconsistent with bulk human RNA-seq. We excluded: (i) 1,225 non-human samples; (ii) 67,443 samples annoated as non-bulk RNA-seq; (iii) 124 in situ assays; and (iv) 22,555 samples with perturbation inconsistencies (defined as samples annotated as “Perturbed” in GSM_Pert but lacking corresponding perturbation info). These steps left 627,482 samples remaining. We then applied study-level and annotation-quality curation. This included removing 19,669 GSM-GSE pairs from GSEs containing fewer than six unique GSMs, excluding 74 samples explicitly labeled as mixed sex, and removing records with composite or overly broad genetic disease terms, non-disease descriptors, or vague or malformed tissue and age annotations. The resulting GEOMeta dataset comprised 594,989 curated GSM–GSE records, corresponding to 594,304 unique GSM accessions from 22,782 GEO Series across diverse experimental studies and biological conditions. Among the final release, 473,770 GSM-GSE records had matched processed expression profiles available through ARCHS4. A schematic overview of the retrieval and filtering workflow is provided in Supplementary Fig. S4.

### Multi-Stage Annotation Pipeline

To address the complexity and heterogeneity of GEO metadata, we implemented an automated annotation framework utilizing a multi-stage agent architecture. The framework uses both study-level descriptions (GSE information) and sample-level descriptions (GSM information) to generate one structured annotation record per GSM under a predefined schema (Supplementary Table S7). The workflow consists of metadata retrieval, role-based extraction, extraction-level QA, field-level standardization, controlled inference, ontology/resource mapping, and deterministic release QA. The overall annotation workflow is shown in Figure 1B. In GEOMeta, an agent refers to an LLM invocation configured with task-specific instructions, a defined metadata context, restricted target fields, and a structured output contract. The agent-based components comprise four task-specific annotation agents in Stage 1, 21 field-specific standardization agent tasks and three controlled inference agent tasks in Stage 2, and four mapping agents in Stage 3. QA1, QA2, and Stage 4 are implemented as deterministic validation procedures, whereas QA3 provides optional evidence-grounded verification and targeted rescue for selected issues. All outputs are required to follow a predefined schema, enabling programmatic validation of structural consistency across batches. Outputs failing schema-level checks, including malformed JSON, incorrect GSM ordering, or row-count mismatches, undergo automated recovery procedures, including constrained reformatting, JSON reconstruction, and row-preserving repair. Expected GSM counts are tracked for each batch to minimize silent sample loss during annotation. Intermediate outputs and audit records are retained throughout the workflow to support reproducibility, targeted review, error tracing, and stage-specific reprocessing. The released pipeline uses a configurable OpenAI-compatible LLM interface, allowing the model, API endpoint, and API key to be specified without modifying the core workflow. Model-specific settings used in individual analyses are reported separately. GPT-5 was used for the annotation, standardization, and mapping experiments reported in this study.

#### (1) Metadata Retrieval and Preparation (Stage 0)

The workflow begins with a user-provided list of GEO Series accessions. In Stage 0, the associated GSM samples are retrieved together with their study-level and sample-level metadata and organized into GSE-linked metadata records. Study-level context is retained for all GSMs from the same GSE. Small GSEs are kept together, whereas larger GSEs are partitioned only when required by the metadata token budget. Partitioning occurs between complete GSM metadata blocks so that individual sample records are not truncated. This study-aware partitioning preserves the shared GSE context while allowing large studies to be processed within model input limits. GEO responses are cached locally, and retrieval failures are retried and recorded in run-specific ledgers and review files. Retrieved metadata and prepared Stage 0 inputs can be reused across model runs, avoiding repeated metadata retrieval and extraction when the same cohort is evaluated using multiple LLM backends.

#### (2) Task-Specific Annotation Agents (Stage 1)

In Stage 1, structured metadata fields were extracted from GEO records using a four task-specific agent annotation strategy. Instead of extracting all metadata fields simultaneously, the task was divided among four agents responsible for related groups of fields, including biological context, experimental context, perturbation, and sample metadata. Each agent used the same GSE-level and GSM-level inputs but applied different field-specific instructions, target-field restrictions, and output constraints.

The biological context agent focused primarily on disease annotation and used a conservative extraction strategy restricted to explicitly supported disease information. The experimental context agent extracted technical and study-level attributes such as sequencing type, RNA library preparation, experimental setting, RNA source, and model type. The perturbation agent extracted perturbation status and associated attributes including perturbation name, dose, duration, frequency, and route of administration, while preserving consistent ordering across multi-condition experiments. The sample metadata agent extracted subject-level and sample-level descriptors such as age, sex, race, ethnicity, sample type, specimen type, timepoint, and outcome. A summary of agent assignments and output-field scope is provided in Supplementary Table S8.

This design reflects how information is organized in GEO metadata. Relevant annotations are often distributed across study descriptions, sample characteristics, titles, and free-text fields rather than appearing in a single structured location. Extracting all fields simultaneously increased the likelihood of omissions, field drift, and cross-field inconsistencies. Separating the task into task-specific agents helped reduce these issues while preserving shared study context across annotations. For each metadata chunk, the four agents received the same GSE-level and GSM-level context and returned agent-specific structured outputs. To reduce runtime, up to two agent requests were processed concurrently, with a short delay between request starts. Call-level failures, including malformed JSON and transient API errors, were retried under a bounded policy allowing up to three total attempts by default. Outputs with row-count or ordering discrepancies underwent row-preserving recovery. The resulting outputs were merged into one structured annotation record per GSM-GSE pair. When the source metadata did not support a value, the corresponding field remained NA rather than being assigned an unsupported value. Agent-call failures that persisted after all permitted attempts were recorded for review and replaced with NA values for the affected fields. Full agent instructions, output schemas, and implementation details are available in [the project GitHub repository](https://github.com/Bin-Chen-Lab/GEOMeta/tree/main/mappings).

#### (3) Extraction-Level Quality Control and Targeted Correction

After raw Stage 1 extraction, a three-part quality Control (QA) stack was applied. This step was designed to detect extraction errors early, because information missed or incorrectly assigned during extraction cannot always be recovered during downstream normalization or ontology mapping.

QA1 performed deterministic within-GSE consistency auditing. For fields expected to remain stable within a study, the procedure summarized missingness and value distributions across all GSMs in each GSE. Missing cells were filled automatically only for eligible fields when all non-missing values within the GSE agreed and field-specific minimum-support and maximum-missingness thresholds were satisfied. Existing non-empty annotations were not overwritten. Fields that may legitimately vary across experimental groups, including age, sex and sample-level perturbation details, were audited but were not corrected using the general within-GSE propagation rule. Ambiguous, conflicting or mixed-design cases were retained for review.

QA2 performed deterministic cross-agent and cross-field validation after the four agents outputs were merged. It evaluated whether related annotations were mutually consistent. For example, QA2 checked whether sample-level perturbation status agreed with the reported perturbation name and, when available, its dose and duration; whether tissue and RNA source annotations were compatible; and whether disease or experimental-setting labels were consistent with the sample type and specimen context. QA2 generated an issue report for subsequent QA3 review but did not directly overwrite Stage 1 annotations.

QA3 performed evidence-grounded verification and targeted rescue using the original metadata. It supported three operating modes: off, which skipped QA3; smart, which reviewed selected high-priority issues identified by QA1 and QA2; and full, which performed a broader QA3 audit. In smart mode, the analysis focused on study-level perturbation status, sample-level perturbation status, RNA source and tissue. Only the flagged field was re-evaluated rather than reannotating all fields for all samples. Corrections were applied automatically only when they were supported by the source metadata and reached the predefined confidence threshold of 0.90. QA3 filled missing values without overwriting existing non-empty annotations. Unsupported, conflicting or low-confidence recommendations were retained in the human-review queue. Due to QA3 is an optional targeted-rescue component, the mode used for each analysis is reported with the corresponding model and pipeline settings.

#### (4) Field-Level Standardization Agent Tasks (Stage 2)

After extraction and targeted correction, field-level standardization was applied to reduce variability in extracted metadata values and improve consistency across studies. This stage was implemented through 21 field-specific standardization agent tasks to harmonize free-text terminology, units, and formats. Standardization procedures included abbreviation expansion, synonym consolidation, capitalization and formatting normalization, and removal of non-informative modifiers or identifiers. Standardization was performed on unique values. For each field, distinct extracted values were collected, normalized into a consistent representation, and then propagated back to all samples containing the corresponding original value. This approach improved computational efficiency and ensured that repeated terms were handled consistently across the dataset. For key metadata fields, both the original raw extracted values (Field_Pre) and the post-standardized outputs (Field_Post) were retained in the full annotation outputs to support traceability. The agent instructions for the 21 field-specific standardization tasks are provided as Markdown files in the GEOMeta GitHub repository. Additional standardization procedures for individual metadata domains are described in the Supplementary Text 1. Tissue annotations were normalized through rule-based harmonization of anatomical and tissue descriptors, with selected brain subregion resolution retained where biologically meaningful (Supplementary Text 1.1). Age annotations were normalized into standardized numeric-unit formats when possible (Supplementary Text 1.2; Supplementary Table S9). Disease annotations underwent multi-stage normalization including deterministic synonym reduction, similarity-based candidate retrieval, constrained model-assisted selection, and manual review (Supplementary Text 1.3). Perturbation-related fields were standardized separately to harmonize treatment names, dose units, duration formats, frequency descriptions, and route-of-administration labels across studies (Supplementary Text 1.4). CP terms were further standardized and mapped to PubChem compound records and structural descriptors (Supplementary Text 1.5). RNA source annotations were standardized using field-specific instructions to harmonize source terminology and formatting (Supplementary Text 1.6).

#### (5) Controlled Inference and Derived Annotations

In addition to direct extraction and standardization, Stage 2 included three task-specific controlled inference agent tasks for age-group derivation, perturbation-type classification, and restricted sex inference from anatomical context. Each task used dedicated instructions and constrained output categories. These derived annotations were maintained separately from directly extracted metadata values.

Sex was first extracted directly from the GEO metadata in Stage 1. Sex inference was applied only when sex information was missing or unclear, anatomical context was considered only for clearly sex-specific tissues such as prostate, testis, ovary, uterus, or placenta. The directly extracted sex annotation and the tissue-derived inference were then merged before sex standardization, with the directly extracted value taking precedence and tissue-based inference used only when the extracted value was missing. The merged values were subsequently standardized to generate the final Stage 2 sex annotation. Neutral or embryonic anatomical terms did not support sex inference. Some secondary labels were generated through controlled derivation steps applied after standardization. Age group annotations were derived from standardized age values using predefined age-bin rules. Numeric ages were assigned to categories such as Pediatric, Adolescent, Adults-20s, Adults-30s, Adults-40s, Adults-50s, and Elderly subgroups. Fetal, newborn, and infant samples were assigned to Infant categories, while cell lines, organoids, experimental timing values, and missing ages were assigned NA (Supplementary Table S9). This derivation step was applied only when the age value represented the biological or developmental age of the human biospecimen.

Perturbation type was similarly derived from normalized perturbation labels using a fixed set of categories including Control/Untreated (CTL), Chemical Perturbation (CP), Biological Perturbation (BIO), Knockout (KO), Knockdown (KD), Overexpression (OE), Environmental Stress (ES), Viral Infection (VIR), OTHER, and NA. Untreated or vehicle-control conditions were assigned CTL, small molecules and drugs were assigned CP, cytokines and antibodies were assigned BIO, and viral infection conditions were assigned VIR, and annotations not covered by the predefined categories were assigned NA, as appropriate. Combination perturbations were allowed to retain multiple perturbation category codes while preserving the original perturbation order.

These derivation and inference steps were intentionally limited to fields with explicit rules and constrained output categories. The purpose was not to infer unsupported biological attributes, but to generate standardized secondary labels from already extracted or normalized metadata values.

#### (6) Ontology Mapping Agents and External Resource Integration (Stage 3)

After standardization and derived-label generation, standardized annotations were linked to external ontology and reference resources. Tissue annotations were mapped to Human Protein Atlas-derived tissue categories (Supplementary Text 1.1), disease annotations were mapped to CTD MEDIC terms (Supplementary Text 1.3), CP names were mapped to CTD chemical terms and PubChem identifiers and descriptors where applicable (Supplementary Text 1.5), and RNA source annotations were mapped to reviewed RNA-source categories and cell-line reference records (Supplementary Text 1.6). These operations were implemented through four task-specific mapping agents for disease, tissue, RNA source, and chemical perturbations.

Ontology mapping followed a reuse-first strategy. Previously curated mapping tables were applied before new mapping attempts were performed. These reusable mapping resources were generated during earlier curation rounds using the same workflow, including field-level standardization, ontology-based candidate retrieval, constrained model-assisted selection, and manual review. Candidate mappings were evaluated using the original sample- and study-level metadata context together with ontology records before incorporation into persistent mapping resources. Previously unseen terms were processed using constrained candidate retrieval and model-assisted selection where applicable. Results that could not be confidently resolved, or that required confirmation before reuse, were retained in field-specific review queues. Only validated mappings were subsequently incorporated into the reusable mapping tables. Tissue annotations that could not be directly mapped to the controlled vocabulary were placed into a tissue review queue. For example, labels such as “Skin: Tumor”, “Skin: Healthy”, or “Blood: Peripheral” contain biologically meaningful information but do not necessarily correspond to existing ontology entries. These terms were reviewed using both sample-level and study-level context to determine whether they could be mapped to existing ontology concepts, incorporated into the curated mapping resources, or retained for further review.

This review-queue framework was also applied to disease and perturbation annotations. Previously unseen terms were retained for manual evaluation and ontology refinement rather than being forced into potentially incorrect mappings. Chemical perturbations were mapped to reviewed compound mappings and PubChem records where applicable. The chemical-perturbation mapping agent generated structured query terms for unresolved single-compound chemical perturbations after reuse of reviewed mappings and cached query results. To limit request frequency, the implementation enforced at least 0.30 s elapsed between consecutive PubChem requests. Temporary server, timeout and throttling responses were handled using status-aware waiting and retry logic, and both successful and unsuccessful query results were cached to avoid repeated requests for the same normalized compound term. PubChem responses were accepted only when the returned compound name or synonym supported the queried perturbation term; ambiguous or unverified results were retained for review. Through iterative review and incorporation of validated mappings into the reusable mapping resources, ontology coverage improved over time while maintaining consistency across annotation runs and dataset releases. This strategy reduced redundant processing, limited ontology drift, and improved reproducibility across studies.

After disease ontology mapping, each disease annotation was further assigned to a curated higher-level disease category representing the dominant clinical or biological disease context. Categories included Oncology, Immune, Infection, Cardiovascular, Respiratory, Digestive, Genetic, Neurology, Metabolic, Congenital, Dermatology, Psychiatry, Musculoskeletal, Reproductive/Fertility, Genitourinary/Urogenital, Pregnancy complications, and Signs/symptoms. The overall grouping framework was adapted from prior therapeutic-area categorization strategies used in large-scale disease analyses [36]. Assignment was based primarily on the dominant disease mechanism or clinical context rather than affected anatomical location. For example, infectious diseases were grouped under Infection even when involving specific organs, while inherited syndromes were grouped under Genetic unless the primary disease identity was neoplastic. Terms describing pathological findings, clinical observations, or non-specific abnormalities were grouped under Signs/symptoms. Category assignment combined rule-based mapping, iterative review, and manual refinement to improve consistency across related disease groups.

Perturbation dose and duration were first standardized in Stage 2 using field-specific post-processing procedures. During Stage 3 release mapping, a dose was retained as a standardized value only when it represented a single interpretable amount with a supported unit, including molar concentrations, mass-per-volume concentrations, body-weight-normalized doses and selected absolute mass units. A duration was retained only when it represented a single value in minutes, hours, days, weeks or months. Multiple values, ranges and other non-canonical expressions were assigned Others, whereas missing dose or duration information was assigned NA. Together, these stages transformed heterogeneous free-text GEO metadata into structured, traceable, and ontology-linked annotations suitable for large-scale transcriptomic analysis.

#### (7) Deterministic Release Quality Assurance (Stage 4)

After ontology and resource mapping, a deterministic release QA stage 4 was applied to evaluate mapped annotations before public release. This step used rule-based validation against controlled vocabularies and consistency constraints. Disease mappings were checked to ensure that final mapped disease labels corresponded to CTD MEDIC disease names or approved non-disease labels, including Normal, Adjacent Normal, No Disease Mentioned, or NA. Disease-associated labels were also required to retain corresponding disease identifiers where applicable. Tissue mappings were checked against the approved tissue vocabulary, including canonical tissue categories and approved brain subregions. Broad disease categories were checked against the predefined category set.

Stage 4 also evaluated consistency across related annotations. The same standardized disease term was required to map to a consistent final disease label and broad disease category. Cancer-, carcinoma-, tumor-, and neoplasm-like terms were flagged if they were not assigned to Oncology. Global and within-GSE consistency checks were applied to key field families, including RNA library, RNA source, disease, tissue, perturbation, age, and sex, using retained Pre, Post, and Mapped values. CP rows were checked for PubChem mapping completeness, including PubChem name, CID, and canonical SMILES for CP rows, while control or non-CP rows were required to have non-applicable PubChem fields. A CP Mapping Status field was retained in internal and CP-focused outputs to distinguish mapped CP rows, non-applicable non-CP/control rows, and CP rows requiring mapping review.

Rows failing deterministic release QA were flagged in internal audit fields and review workbooks rather than silently included as release-ready annotations. Public release files excluded detailed QA columns but retained curated biological annotations, disease identifiers, PubChem fields where applicable, and provenance metadata. The filtered mapped outputs, Stage 4 QA reports, and review queues were retained separately to support auditability and targeted expert review.

To improve reproducibility and targeted reprocessing, all intermediate outputs are retained throughout the pipeline rather than overwritten. Structured outputs include tabular annotation files, row-level structured records, retrieval and chunk-processing ledgers, schema-validation summaries, failed-sample reports, Stage 1 QA reports, Stage 2 review and reannotation queues, post-mapping correction queues, Stage 4 release-QA workbooks, and reusable mapping resources. This design enables individual stages or metadata domains to be reprocessed independently without rerunning the full annotation workflow and provides an auditable trace of transformations applied to each sample.

### Evaluation Design

Because GEO lacks comprehensive gold-standard annotations across most metadata domains, evaluation of GEOMeta required multiple complementary validation strategies rather than reliance on a single reference dataset. Annotation quality control therefore relied on layered mechanisms throughout the workflow, including role-based context-aware extraction, schema-level validation, field-specific semantic normalization, constrained ontology mapping, reusable curated mapping resources, novel-term review queues, targeted manual review, benchmark-based evaluation using expert-reviewed samples, external concordance analyses, and text-evidence validation against original GEO metadata. Together, these components assessed annotation consistency, biological plausibility, robustness to prompt design, ontology-aware agreement, and reproducibility across heterogeneous metadata fields while preserving auditability throughout the annotation process. The evaluation cohorts used in this study are summarized in Supplementary Table S10.

#### (1) Benchmark Dataset for Annotation Evaluation

To support prompt development and assess annotation consistency and support prompt refinement, we constructed a benchmark dataset of 1,000 GEO samples through manual curation. These samples were generated during the prompt optimization process by reviewing and correcting outputs from the annotation pipeline, followed by independent validation by domain experts using the original GSE- and GSM-level metadata. This benchmark was developed iteratively alongside prompt refinement. During this process, annotation rules and field definitions were updated based on expert feedback, particularly for fields requiring contextual interpretation or complex normalization. We focused the benchmark evaluation on key fields included in the final released dataset. Ultimately, the benchmark dataset included 1,000 samples and five fields: sex, experimental setting, age, disease, and tissue. For these fields, agreement between automated pipeline annotations and manual curation was calculated as the proportion of matching annotations. This comparison reflects performance at the level of extracted and lightly standardized values, prior to full ontology mapping. Because the benchmark dataset was developed iteratively during prompt refinement, this analysis is intended to assess consistency with expert-reviewed annotations. Prompts were updated based on observed errors, expert feedback, and recurring edge cases, and were then re-evaluated on the same benchmark set. Over successive iterations, performance stabilized across evaluated fields. In later iterations, changes to prompt wording did not consistently improve performance across samples. Instead, they often resolved some edge cases while introducing new inconsistencies in others, suggesting that the prompts had reached a stable configuration. This benchmark therefore serves as an internal validation framework for prompt design, ensuring that the final annotation pipeline achieves consistent performance on core fields before large-scale deployment.

#### (2) Prompt Design Ablation

To evaluate prompt design, we constructed a separate 1,000-sample comparison dataset by randomly selecting 200 GSEs and five GSM samples from each GSE. This dataset was not used for prompt development or refinement. The same samples were annotated using the task-specific Stage 1 agent prompts and a generic structured prompt, while Stage 2 standardization and Stage 3 mapping were held constant. Comparisons included RNA library, experimental setting, RNA source, disease, tissue, age, age group, sex, GSE-level perturbation, GSM-level perturbation and perturbation identity. For each field, we calculated missingness under each prompt configuration, one-sided non-missing outputs, discordant outputs among samples annotated by both prompts and concordance among shared outputs. Stage 1 and final output differences were defined as the sum of task-specific-only, generic-only and discordant outputs across all 1,000 samples. Untreated and explicit control labels were treated as valid perturbation values. Neither prompt configuration was treated as a reference standard; therefore, the comparison assessed annotation availability and cross-prompt agreement rather than accuracy.

#### (3) External Validation Datasets

GEOMeta annotations were externally evaluated using two independently curated resources. DiSignAtlas [37] was used to assess disease and tissue annotations, whereas PharmGEO [38] was used to assess CP annotations. Neither resource was used during GEOMeta prompt development, annotation generation, or mapping-resource construction. Records were compared with GEOMeta at the GSM level after resource-specific identifier harmonization and preprocessing. DiSignAtlas preprocessing and matching are described in Supplementary Text 2.1, whereas PharmGEO processing and drug-name reconciliation are described in Supplementary Text 2.2.

#### (4) Text-Evidence Validation for Sex and Age

Because no independent external reference dataset was available for sex and age, we performed a text-evidence consistency analysis using the original GEO metadata. This analysis was designed to evaluate the initial extraction step. Therefore, text-derived values were compared with the raw GEOMeta extraction outputs for Sex and Age, rather than the standardized outputs. Post-processed fields were not used because they include additional normalization and controlled inference, while the goal here was to assess whether Stage 1 captured explicitly reported sex and age information from the metadata. Rule-based extraction considered both GSM-level and GSE-level metadata. However, because sex and age are sample-level attributes, GSM-level metadata were prioritized. GSE-level metadata were used only when the information appeared in an explicit field-like form and could reasonably be interpreted as sample-specific. General cohort descriptions, eligibility criteria, or study-level summaries were not used to assign sex or age to individual samples.

For sex, the rule-based extractor captured explicit entries such as “Sex: Male,” “Sex: F,” “Gender: Female,” or “biological sex: male.” These values were normalized to Male, Female, or Mixed. Broad mentions of males or females in study descriptions were excluded to avoid assigning group-level statements to individual samples.

For age, the extractor captured explicit age-related fields from the metadata, including numeric values with units such as years, months, weeks, or days. Age values were normalized into a consistent numeric-unit format, with standardized capitalization and rounding of decimal values to two places. Common unit abbreviations and developmental formats were also normalized before comparison. However, because GEO age fields are frequently ambiguous, the extracted text-derived age values were manually reviewed before comparison. This review step was necessary because numeric age entries may represent developmental stage, experimental duration, passage number, treatment timing, or non-human host age rather than the biological age of the human biospecimen. During manual review, incorrect or contextually inconsistent rule-based age interpretations were corrected using the original GSE and GSM metadata prior to comparison. Broad cohort descriptions and eligibility criteria were not used to infer sample-level age values.

Before comparison, missing values, including “NA,” “Unknown,” “not reported,” and equivalent labels, were converted to empty values. For each field, we summarized four categories: both missing, both non-missing with matching values, both non-missing with mismatching values, and one-sided missingness. Shared missingness was analyzed separately and was not counted as agreement. Agreement was calculated only among samples where both GEOMeta extraction and text-derived extraction produced non-missing values.

#### (5) Independent Evaluation Cohort and Expert Review

To evaluate GEOMeta on recently released GEO records not used during pipeline development, we constructed an independent May 2026 evaluation cohort. We queried the NCBI GEO DataSets database for GEO Series records with a publication date in May 2026. The cohort was defined at the GSE level rather than the GSM level, so all samples associated with an eligible GSE were retained together. We restricted the cohort to strict human-only studies by retaining GSE records whose GEO DataSets taxon field exactly matched Homo sapiens, thereby excluding mixed-organism records. GSEs were also excluded if the associated GSM publication month differed from the GSE publication month. We applied a data-driven GSE-size filter based on the May 2026 human-only GEO distribution, retaining GSEs with 6-600 associated GSM samples. This range excluded many very small studies and a small number of large outlier studies that would disproportionately affect runtime, cost, and sample burden. Eligible GSEs were then selected using stratified sampling across GSM-count bins: 6-20, 21-50, 51-100, 101-200, 201-400, and 401-600 GSMs per GSE. This provided a heterogeneous recent-release evaluation set with a broad distribution of study sizes. The final selected cohort was processed through the GEOMeta pipeline using GPT-5. For expert evaluation, public-style release files were shared with a domain expert in pharmacology and toxicology. The expert reviewed selected randomized rows and, when needed, examined entire studies to verify difficult or potentially inconsistent annotations. Manual review focused on disease, tissue, RNA source, age, and sex fields. Expert comments were compared against GEOMeta-generated annotations to identify confirmed annotations, non-confirmed annotations, and recurring sources of error.

#### (6) Multi-LLM benchmarking and prospective annotation

Before the larger prospective analysis, we evaluated 22 LLMs from 11 model providers available through OpenRouter using an internal benchmark cohort of 118 samples from five GEO Series submitted by our laboratory. The benchmark was used only for model comparison and selection. Each model was evaluated using the same metadata inputs and was processed using the same GEOMeta workflow. Benchmark consensus annotations were constructed from models producing the expected complete release outputs. For the final-release benchmark, 16 models retaining all 118 samples were used to construct the consensus annotations. For the CP benchmark, nine models whose CP-release files contained the expected 18 benchmark samples were used to construct a separate CP consensus. Model outputs were aligned by GSM identifier and compared after applying the same value-normalization procedures used in the GEOMeta pipeline. For each evaluable field, the most frequently supported normalized value, including the literal category “NA,” was selected. Ties for the highest support were treated as unresolved, and the corresponding fields were excluded from the agreement evaluation.

All 22 model outputs were subsequently compared with the corresponding benchmark consensus annotations. Final-release output completeness was defined as the proportion of the 118 benchmark samples retained in each model-specific final-release file, and CP output completeness was defined relative to the 18 CP benchmark samples. A benchmark sample absent from a model-specific release file was recorded as no prediction and counted as discordant in the completeness-penalized agreement metric. The literal annotation value “NA” was treated as a valid returned category and was considered concordant only when the consensus annotation was also “NA.” Final-release agreement was evaluated across 15 core annotation fields, and CP agreement was evaluated across eight perturbation-specific fields.

For each model, we recorded final-release output completeness, concordance with the final-release consensus, CP output completeness, concordance with the CP consensus, runtime and API cost. Model selection first considered output completeness and consensus concordance. Runtime and API cost were then considered among models producing sufficiently complete and concordant outputs. This procedure resulted in the selection of Gemini 3 Flash Preview, Mistral Large 2512, GPT-5 mini, DeepSeek V4 Flash and Qwen3.7 Plus.

The May 2026 cohort was used for independent expert review. Because only a limited number of May 2026 GSEs had matched expression profiles in the ARCHS4 version used for downstream analysis, we selected a separate expression-linked cohort comprising GEO Series submitted in December 2025 and January 2026. These records were excluded from the initial GEOMeta release. The selected GSEs were matched against sample metadata in the ARCHS4 human gene-level expression file to identify samples with available expression profiles. Studies with fewer than six associated samples were excluded. Parent SuperSeries were also excluded to avoid including the same samples through both parent and component GSE records. Each of the five selected models was applied independently to this same set of GSEs using the same pipeline configuration and metadata inputs. Runtime was recorded as the end-to-end processing time, and API cost was recorded.

Consensus annotations were constructed at the GSM level across the five model outputs. Independent fields were compared after value normalization. A model that did not contain a given GSM contributed no vote for that sample, whereas blank field values within retained GSM records were normalized to “NA” and included in voting. Ties for the highest support were treated as unresolved, and samples with an unresolved tie in an independent or primary field were excluded. Disease was used as the primary disease field, and DiseaseID, AltDiseaseIDs and Broad_Disease_Category were taken from models supporting the selected Disease value. Age was used as the primary age field, and Age_Group was taken from models supporting the selected Age value. GSM_Pert was determined at the sample level. GSE_Pert was then derived from the retained GSM_Pert annotations within each GSE: GSE_Pert was assigned as Yes if at least one retained sample was perturbed and as No if no retained sample was perturbed. Perturbation was treated as the primary perturbation field in the CP-focused output, and Perturbation Type, Perturbation Dose, Perturbation Duration and PubChem-related fields were taken from models supporting the selected Perturbation value. Samples with unresolved ties in an independent or primary field were excluded from the clean consensus release. The final CP release was further restricted to GSEs containing both control-like and perturbed retained samples.

#### (7) Dataset Description and Data Splits

The final GEOMeta dataset includes large-scale sample-level annotations derived from GEO metadata, with standardized and mapped fields covering study identifiers (GSE_ID), sample identifiers (GSM_ID), study year, contact country, sequencing platform, RNA library, experimental setting, perturbation status, disease, disease category, tissue, sex, and age group. In addition to the full annotated dataset, we provide predefined training, study-held-out test set, and prospective test (“2024 temporal holdout set”) splits to support downstream benchmarking and model development.

In addition to serving as a large-scale annotated resource, the dataset is structured to support several downstream tasks. First, GEOMeta enables systematic cohort construction for drug perturbation studies by identifying treated and control samples across GEO, supporting cross-study drug signature discovery and meta-analysis. Second, standardized disease and normal annotations allow disease-specific cohorts to be assembled across studies, supporting disease signature discovery and drug–disease comparison analyses. Third, standardized labels for disease, tissue, age group, and sex provide benchmark tasks for evaluating transcriptome-based prediction models. The predefined train, study-held-out test set, and prospective test sets further support consistent evaluation of both in-distribution performance and generalization to newer studies.

#### (8) Downstream Benchmarking for Label Prediction

We established a benchmarking framework to evaluate how well current transcriptome representation models capture major biological features. Four tasks: Sex, Tissue, Disease, and Age Group were used, as they represent broad biological variation and are commonly used to assess model utility.

To avoid study leakage, data splitting was performed at the GSE level. The training and study-held-out test set were constructed from the full dataset, with 10% of GSEs held out for testing. Only labels present in the training set were retained in the test set. In addition, we used a prospective test set (“2024 temporal holdout set”) consisting of more recent samples not included in the original training distribution. The study-held-out test set was used to evaluate in-distribution performance, while 2024 holdout was used to assess generalization to newer and potentially more heterogeneous data. Both test sets were evaluated and reported separately.

For the age classification task, labels were merged prior to evaluation as follows: Infant, Pediatric, and Adolescent were grouped as “Youth”; Adults-20s, Adults-30s, and Adults-40s as “Adult”; and Adults-50s, Elderly-1, Elderly-2, and Elderly-3 as “Late Adult/Elderly”. To generate the AnnData object for benchmark analyses, transcript abundance profiles were obtained from the ARCHS4 human reference HDF5 file (*human_gene_v2.5.h5*) by reading genes in batches (1,000 genes per chunk) to reduce memory usage, while excluding two known problematic gene index ranges (56,000–58,000). Gene identifiers and sample annotations were extracted from the same HDF5 file and assembled into an AnnData object. Finally, the curated metadata tables were merged by GSM accession to add standardized attributes (sex, organ system, disease, age group, and experimental setting), and duplicate GSM entries were removed.

#### (9) Baseline Evaluation Protocols

We benchmark pretrained single-cell foundation models on bulk RNA-seq samples under two standard transfer learning protocols: MLP probing and full fine-tuning^31,35^. Bulk RNA-seq samples are treated as pseudo-cells, enabling us to assess how well representations learned from single-cell data transfer to bulk settings. All experiments are conducted on the GEOMeta dataset across four downstream prediction tasks: Tissue, Disease, Age, and Sex. Accuracy serves as the primary evaluation metric.

### Problem Formulation

Let *x_i_* ∈ *R^G^* denote the gene expression vector of sample *i*, where *G* is the number of genes. For each method, we obtain a sample embedding *z_i_* = *f_θ_*(*x_i_*), where *f_θ_*(·) denotes the pretrained encoder or latent representation function. Under the probing protocol, the parameters of *f* remain frozen, and only a downstream classifier is optimized. Under the fine-tuning protocol, both *f* and the classification head are jointly updated end-to-end.

### MLP Probing

Given a frozen embedding *z_i_*, we train a two-layer multilayer perceptron (MLP) classifier *h_i_* = σ(*W_1_z_1_* + *b_1_*) *ŷ* = *soft*max (*W_2_h_i_* + *b_2_*), where *h_i_* ϵ *R*^256^ is the hidden representation, *σ*(⋅) is the ReLU activation, and *y*4_&_ is the predicted class probability vector. The classifier is trained with cross-entropy loss using the Adam optimizer (learning rate 1 × 10^*+^, batch size 32) for up to 1,000 epochs.

We evaluate the following three encoders *f*: scGPT^10^ is a transformer-based generative foundation model that treats genes as tokens and learns contextualized gene and cell representations autoregressively. We load the pretrained scGPT_human checkpoint and generate sample embeddings via the embed_data function with a batch size of 64.

CellPLM^13^ is a cell-centric transformer that treats cells as tokens and explicitly models inter-cellular relationships rather than focusing solely on gene–gene interactions. We use the 85M-parameter pretrained model (20231027_85M) and extract embeddings through the CellEmbeddingPipeline with a batch size of 4,000.

scVI^49^ is a variational autoencoder that learns batch-invariant latent representations by conditioning on the GSE ID as a batch covariate. Unlike scGPT and CellPLM, scVI is trained from scratch on the target data: we apply library-size normalization and log transformation, select the top 2,000 highly variable genes, and fit a VAE with two encoder layers, a 128-dimensional latent space, and a negative binomial likelihood.

BulkFormer^9^ is a 150M-parameter transformer pretrained on ∼520,000 human bulk transcriptomes that combines a graph convolutional layer over a prior gene co-expression graph with Performer attention, and is the only baseline pretrained on the same modality as our evaluation data. We load the pretrained BulkFormer-147M checkpoint and extract sample embeddings by mean-pooling the final-layer contextualized gene embeddings over the model’s curated informative-gene subset, with a feature-extraction batch size of 16.

### Fine-Tuning

For fine-tuning, we adapt each pretrained foundation model by attaching a feed-forward classification head and training the entire network end-to-end on the GEOMeta training set with cross-entropy loss. This allows the model to capture task-specific structure while retaining the inductive biases of the pretrained representation.

scGPT (fine-tune). We initialize a TransformerModel from the pretrained scGPT_human checkpoint with an embedding dimension of 128, 4 attention heads, and 4 encoder layers, followed by a 3-layer classifier head with dropout rate 0.2. Gene expression values are binned into 51 bins, and the top 3,000 highly variable genes are retained. The model is optimized with Adam (learning rate 1 × 10^*,^, batch size 16) and a step learning rate scheduler (decay ratio 0.9) for 30 epochs.

CellPLM (fine-tune). We employ the CellTypeAnnotationPipeline initialized from the 85M-parameter pretrained model. Training uses a learning rate of 1 × 10^*+^ or 1 × 10^*,^ for up to 1,000 epochs, with 15% of the training data held out for early stopping. The maximum batch size is 3,000.

BulkFormer (fine-tune). We attach a classification head to the pretrained BulkFormer-147M backbone and train end-to-end with AdamW using separate learning rates for the backbone (1 × 10^*(^) and head (1 × 10^*+^), weight decay (1 × 10^*,^), and linear warmup followed by cosine decay. Training uses a batch size of 16 with 8 gradient-accumulation steps (effective batch size 128) under mixed precision with gradient checkpointing, for 10 epochs.

### Validation of sex-discordant samples using raw RNA-seq data

We reanalyzed raw RNA-seq data from representative samples. Raw FASTQ files were downloaded from the NCBI Sequence Read Archive (SRA) using SRA Toolkit (v3.0.10). Reads were aligned to the human GRCh38 reference genome with STAR (v2.7.11b) using the GENCODE v45 annotation. Alignment parameters retained NH (number of genomic alignments), AS (alignment score), and nM (mismatch count) tags, while allowing up to 50 multimapping locations to explicitly assess mapping ambiguity. Because STAR assigns MAPQ = 255 exclusively to uniquely mapped reads, this metric was used to distinguish uniquely aligned chromosome Y reads from multimapping alignments. Gene-level expression was quantified using both STAR GeneCounts and featureCounts (v2.0.6). For each sample, we summarized total chromosome Y reads, uniquely mapped chromosome Y reads (MAPQ = 255 and NH = 1), multimapping reads (NH > 1), and the fraction of uniquely mapped chromosome Y reads. We further examined expression of canonical sex-marker genes, including *XIST* and multiple Y-linked genes (*RPS4Y1*, *DDX3Y*, *EIF1AY*, *KDM5D*, *UTY*, *USP9Y*, and *ZFY*).

### Dataset Scope, Quality, and Intended Use

GEOMeta was developed as a large-scale standardized metadata resource for human bulk RNA-seq studies derived from GEO and ARCHS4. The dataset is intended to support large-scale transcriptomic analysis, disease signature discovery, drug-response and perturbation analysis, representation learning, benchmark development, metadata prediction, and cross-study integrative analysis. The structured annotations also facilitate downstream applications including biomarker discovery, comparative disease analysis, computational drug repurposing, and transcriptomic foundation model benchmarking. To support reproducible machine learning research and benchmark evaluation, curated metadata tables for the full release and processed AnnData objects for the expression-matched subset were released through the GEOMeta Hugging Face repository.

The released dataset includes study-aware train, test, and held-out evaluation splits together with standardized metadata fields covering disease, tissue, sex, age group, experimental setting, perturbation status, sequencing platform, and RNA library type. The expression-matched subset includes study-aware training, test and held-out evaluation splits. Its processed AnnData objects contain transcript abundance matrices linked with sample-level and gene-level annotations, enabling direct integration into downstream machine learning and bioinformatics workflows.

Although GEOMeta incorporates multiple quality-control steps, including extraction, standardization, ontology mapping, validation, and targeted review, the final annotations remain dependent on the completeness and accuracy of the original GEO submissions. Some fields, particularly disease, tissue, perturbation, and experimental setting, may retain residual ambiguity or reduced granularity after standardization. Broad disease categories were designed primarily to support large-scale analysis and benchmarking and therefore may not preserve all disease-specific clinical distinctions.

The current pipeline prioritizes annotation quality and consistent interpretation over maximum speed. Stage 1 is the main runtime bottleneck because it performs context-aware extraction using multiple role-specific prompts for each GSE/GSM metadata batch. This design is slower than a rule-based or single-prompt approach, but it provides clearer field boundaries and supports more consistent downstream standardization and ontology mapping. An important feature of the framework is its iterative refinement capability: rather than treating unmapped terms as permanent failures, unresolved annotations are retained in review queues and incorporated into subsequent ontology refinement cycles after review. This process allows ontology coverage to improve over time while preserving consistency across releases. Future versions could improve runtime through stronger caching, prompt compression, parallel GSE-batch processing, or smaller specialized models for simpler fields.

The dataset was designed primarily for computational analysis at scale rather than for direct clinical interpretation. Users performing disease-specific, clinically sensitive, or small-cohort analyses should consider additional manual review of relevant metadata fields. Despite these limitations, GEOMeta provides one of the largest publicly available structured metadata resources for human bulk RNA-seq studies and establishes a scalable framework for systematic reuse of public transcriptomic data.

## Data and Code Availability

The GEOMeta dataset, including training, test, and prospective test splits, is available at https://huggingface.co/datasets/binchenlab/GEOMeta

The GEOMeta source code, task-specific agent instructions and prompts, multi-stage processing and quality-control procedures, mapping resources, and configurable LLM backend interface are publicly available at https://github.com/Bin-Chen-Lab/GEOMeta

## Author Contribution

X.Z. coordinated the study, designed and implemented the overall GEOMeta multi-stage agent framework, and performed the large-scale annotations and external validation analyses. X.Z. and S.P. co-developed the task-specific agent prompts and worked on metadata standardization and ontology mapping. S.P. performed the benchmark dataset evaluation. J.P. and Y.X. designed and implemented the downstream label-prediction benchmarks, including model training and evaluation using foundation-model embeddings, fine-tuning approaches, and classical baselines. S.M. prepared the ARCHS4-derived datasets. R.S. and D.L. contributed to disease-field prompt development and annotation review. R.N. performed domain-expert review of annotations in the prospective evaluation cohort. S.K. independently tested the multi-stage annotation workflow and performed the chromosome Y audit of sex-discordant samples. Z.P. independently tested the workflow and performed perturbation-focused validation of case/control annotations. X.L. contributed domain expertise to strain-related prompt development and annotation review. B.C. contributed to prompt and pipeline development, the development and review of metadata-field mappings, and supervised the study. All authors contributed to manuscript preparation.

## Acknowledgement

We thank our summer interns Chris Chang, Jenny Qi and Rohit Agarwal for their contributions to manual metadata curation, and Aravind Gurusaran Korukonda for reviewing and standardizing the curated annotations.

