## Supplemental File for "Automating scientific annotations for open transcriptomic profiles via multi-stage agents"

**Supplementary Files**

- Supplementary Table S1. Field-level coverage and missingness for major metadata attributes in the final curated human GEO dataset
- Supplementary Table S2. Effect of Stage 1 prompt design on final annotation completeness, fragmentation, and information loss
- Supplementary Results 1. External validation
  Supplementary Results 1.1. Disease and tissue concordance with DiSignAtlas
  Supplementary Results 1.2. Chemical perturbation concordance with PharmGEO
- Supplementary Figure S1. Benchmarking transcriptomic representation-learning models across tasks and evaluation settings.
- Supplementary Figure S2. The confusion matrix for age prediction result generated by finetuned CellPLM on 2024 holdout
- Supplementary Figure S3. Sample-level expression profiles of sex-linked marker genes in the 2024 holdout cohort, stratified by classification outcome.
- Supplementary Table S3. Expert manual review confirmation rates for selected GEOMeta annotations in the prospective May 2026 GEO cohort
- Supplementary Table S4. Performance, runtime, and cost comparison of 5 Selected LLMs for GEOMeta annotation on perspective analysis
- Supplementary Table S5. Representative GEO age annotation edge cases
- Supplementary Table S6. Representative GEO tissue annotation edge cases
- Supplementary Table S7. Annotation schema and definitions for the 27 standardized metadata fields
- Supplementary Figure S4. Data retrieval, filtering and preprocessing workflow for the GEOMeta dataset
- Data retrieval, filtering and preprocessing workflow for the GEOMeta dataset.
- Supplementary Table S8. Annotation agents and output-field scope
- Supplementary Table S9. Age-group definitions
- Supplementary Table S10. Summary of evaluation cohorts used in this study
- Supplementary Text 1. Field-specific standardization and mapping

Supplementary Text 1.1. Tissue term standardization and controlled-vocabulary mapping
Supplementary Text 1.2. Age annotation standardization and age-group derivation
Supplementary Text 1.3. Disease term curation, standardization, and ontology mapping
Supplementary Text 1.4. Perturbation-related field standardization and controlled inference
Supplementary Text 1.5. Chemical perturbation standardization and PubChem mapping
Supplementary Text 1.6. RNA-source standardization and mapping

- Supplementary Text 2. External validation methods
  Supplementary Text 2.1. DiSignAtlas preprocessing and disease and tissue concordance assessment
  Supplementary Text 2.2. PharmGEO preprocessing and chemical perturbation concordance assessment

**Supplementary Table S1:** Field-level coverage and missingness for major metadata attributes in the final curated human GEO dataset

| Field | Non missing counts | Missing Counts | Coverage (%) |
|---|---|---|---|
| Experimental Setting | 594,989 | 0 | 100.00 |
| GSE_Pert | 594,989 | 0 | 100.00 |
| RNA Library | 594,760 | 229 | 99.96 |
| RNA Source | 592,201 | 2,788 | 99.53 |
| GSM_Pert | 590,057 | 4,932 | 99.17 |
| Disease | 585,744 | 9,245 | 98.45 |
| Tissue | 519,779 | 75,210 | 87.36 |
| DiseaseID | 315,181 | 279,808 | 52.97 |
| Broad Disease Category | 315,163 | 279,826 | 52.97 |
| Sex | 188,937 | 406,052 | 31.75 |
| Age Group | 96,456 | 498,533 | 16.21 |
| Age | 93,559 | 501,430 | 15.72 |

**Supplementary Table S2.** Effect of Stage 1 prompt design on annotation availability and cross-prompt concordance

| Metadata field | Stage 1 output difference, *n* (%) | Final missing, task-specific (%) | Final missing, generic (%) | Final task-specific only / generic-only, *n* | Final discordant shared outputs, *n* | Final output difference, *n* (%) | Final concordance among shared outputs (%) |
|---|---|---|---|---|---|---|---|
| RNA library | 119 (11.9) | 1.0 | 4.7 | 47 / 10 | 62 | 119 (11.9) | 93.4 |
| Experimental setting | 175 (17.5) | 0.0 | 6.5 | 65 / 0 | 110 | 175 (17.5) | 88.2 |
| RNA source | 990 (99.0) | 0.0 | 4.5 | 45 / 0 | 837 | 882 (88.2) | 12.4 |
| Disease | 141 (14.1) | 41.7 | 37.4 | 15 / 58 | 5 | 78 (7.8) | 99.1 |
| Tissue | 501 (50.1) | 20.6 | 53.8 | 350 / 18 | 5 | 373 (37.3) | 98.9 |
| Age | 16 (1.6) | 87.9 | 88.8 | 10 / 1 | 5 | 16 (1.6) | 95.5 |
| Age group | 127 (12.7) | 87.9 | 88.9 | 10 / 0 | 0 | 10 (1.0) | 100.0 |
| Sex | 11 (1.1) | 80.7 | 74.1 | 0 / 66 | 0 | 66 (6.6) | 100.0 |
| GSE-level perturbation | 30 (3.0) | 0.5 | 0.0 | 0 / 5 | 25 | 30 (3.0) | 97.5 |
| GSM-level perturbation | 92 (9.2) | 3.5 | 2.0 | 10 / 25 | 57 | 92 (9.2) | 94.0 |
| Perturbation | 520 (52.0) | 10.0 | 31.8 | 238 / 20 | 276 | 534 (53.4) | 58.3 |

Notes:

(1) N = 1,000 samples for every field.

(2) "Stage 1 output difference" and "Final output difference" were defined as the sum of task-specific-only, generic-only and discordant outputs and were calculated across all samples.

(3) "Final task-specific only / generic only" reports one-sided non-missing final outputs in that order.

(4) "Final discordant shared outputs" includes only samples for which both prompts returned non-missing final outputs but produced different values.

(5) "Final concordance among shared outputs" was calculated only among samples with non-missing final outputs from both prompts.

(6) Untreated and explicit control labels were treated as valid non-missing perturbation values.

(7) Final sex annotations include downstream controlled inference from sex-specific anatomical context.

(8) Neither prompt configuration was treated as a reference standard; these comparisons assess annotation availability and cross-prompt agreement rather than annotation accuracy

### Supplementary Results 1. External validation

**Supplementary Results 1.1.** Disease and tissue concordance with DiSignAtlas

After sample-level expansion, duplicate removal, and matching to GEOMeta, 19,879 samples from 540 GSEs were retained for evaluation. DiSignAtlas preprocessing and sample-level matching procedures are described in Supplementary Text 2.1.

For disease concordance analysis, samples annotated as Normal, Adjacent Normal, or No Disease Mentioned were excluded, leaving 13,075 disease-associated samples from 453 GSEs. Of these, 81.75% showed direct disease agreement, 0.69% showed synonym agreement, and 14.15% were classified as generalized matches, including subtype–parent disease relationships. Overall, 96.59% of disease-associated samples showed concordant annotations, whereas 3.41% remained unmatched.

Disease-status concordance was assessed separately by grouping Normal, Adjacent Normal, and No Disease Mentioned as non-disease and all other samples as disease-associated. Across the 19,879 matched samples, 96.23% showed concordant assignments, with non-disease samples corresponding to DiSignAtlas controls and disease-associated samples corresponding to cases. The remaining 3.77% showed discordant assignments.

For tissue evaluation, 23.64% of the 19,879 matched samples lacked a tissue annotation in at least one resource and were excluded from the paired-label analysis. Among the 15,179 samples with tissue annotations available in both resources, 96.95% showed concordant standardized tissue annotations, whereas 3.05% showed disagreement.

**Supplementary Results 1.2.** Chemical perturbation concordance with PharmGEO

After restricting PharmGEO to human samples, 26,548 unique GSMs were available for integration. Exact matching by GSE and GSM identifiers identified 10,359 samples shared between PharmGEO and GEOMeta, of which 59.3% were treated samples and 40.7% were controls according to PharmGEO.

Initial comparison of compound annotations showed 85.81% concordance, whereas 14.19% were initially classified as discordant. Many apparent discrepancies reflected differences in naming conventions, including abbreviated compound names, salt or formulation variants, partial representation of multi-compound perturbations, and chemical synonyms. After applying a curated equivalence table, 74.22% of the initially discordant samples were reclassified as concordant. Overall concordance therefore increased to 96.34%, with 3.66% remaining discordant. Treatment-status concordance was also high. PharmGEO treated/control assignments agreed with GEOMeta perturbation type for 95.48% of samples and with GEOMeta sample-level perturbation status for 96.44%. Within GEOMeta, agreement between perturbation type and sample-level perturbation status exceeded 98%.

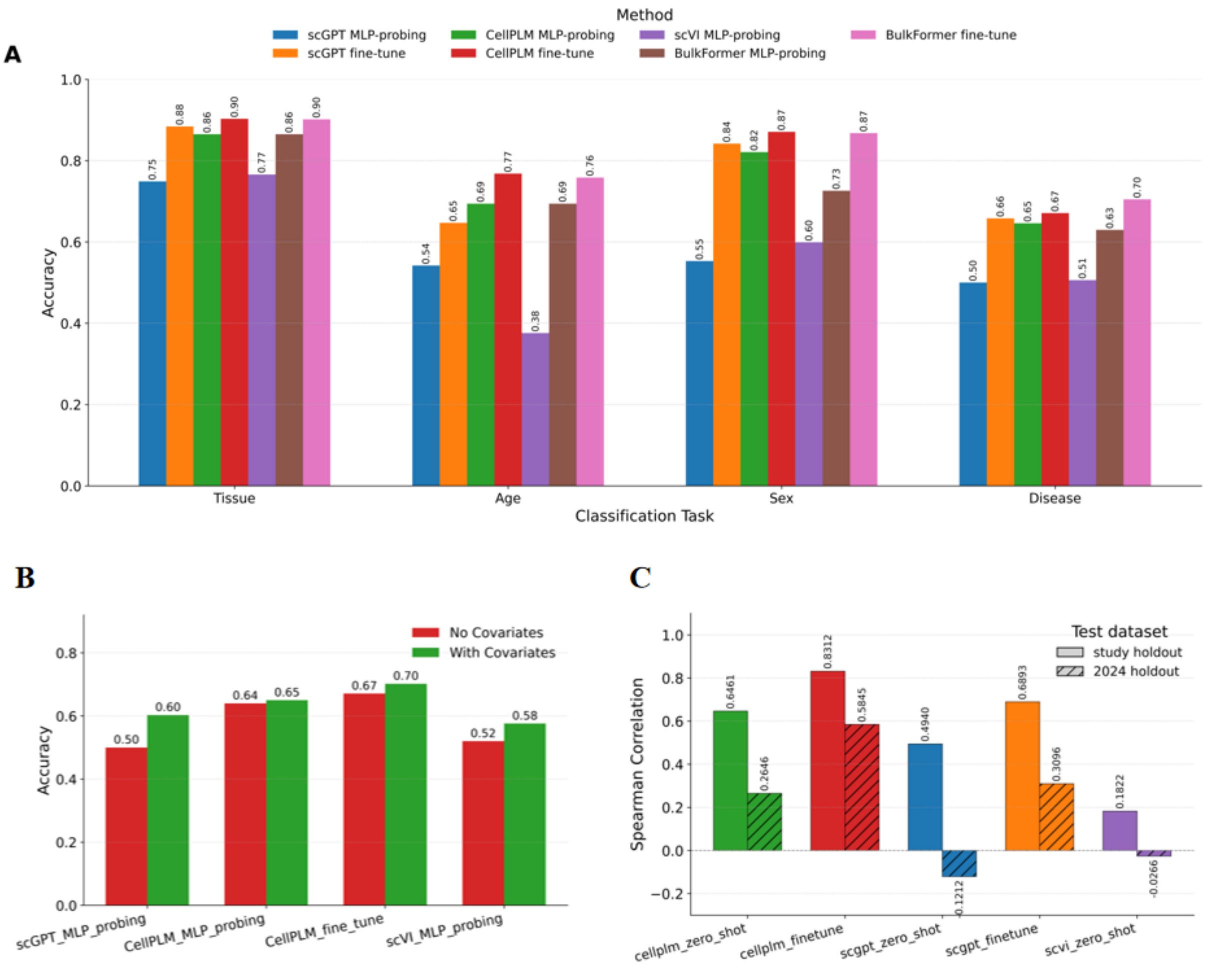


**Supplementary Figure S1. Benchmarking transcriptomic representation-learning models across tasks and evaluation settings. A:** Label prediction results for the study holdout dataset. Accuracy achieved by seven methods scGPT (MLP-probing), scGPT (Fine-Tune), CellPLM (MLP-probing), CellPLM (Fine-Tune), scVI (MLP-probing), Bulkformer (MLP-probing), Bulkformer (Fine-Tune) across four prediction tasks (Sex, Organ, Disease, and Age). **B:** Disease prediction results for the study holdout datasets. Accuracy achieved by four methods scGPT (MLP-probing), CellPLM (MLP-probing), CellPLM (Fine-Tune), scVI (MLP-probing) with and without additional covariate information. **C:** Age prediction results for the study holdout and 2024 holdout datasets. Spearman correlation achieved by five methods scGPT (MLP-probing), scGPT (Fine-Tune), CellPLM (MLP-probing), CellPLM (Fine-Tune), scVI (MLP-probing).

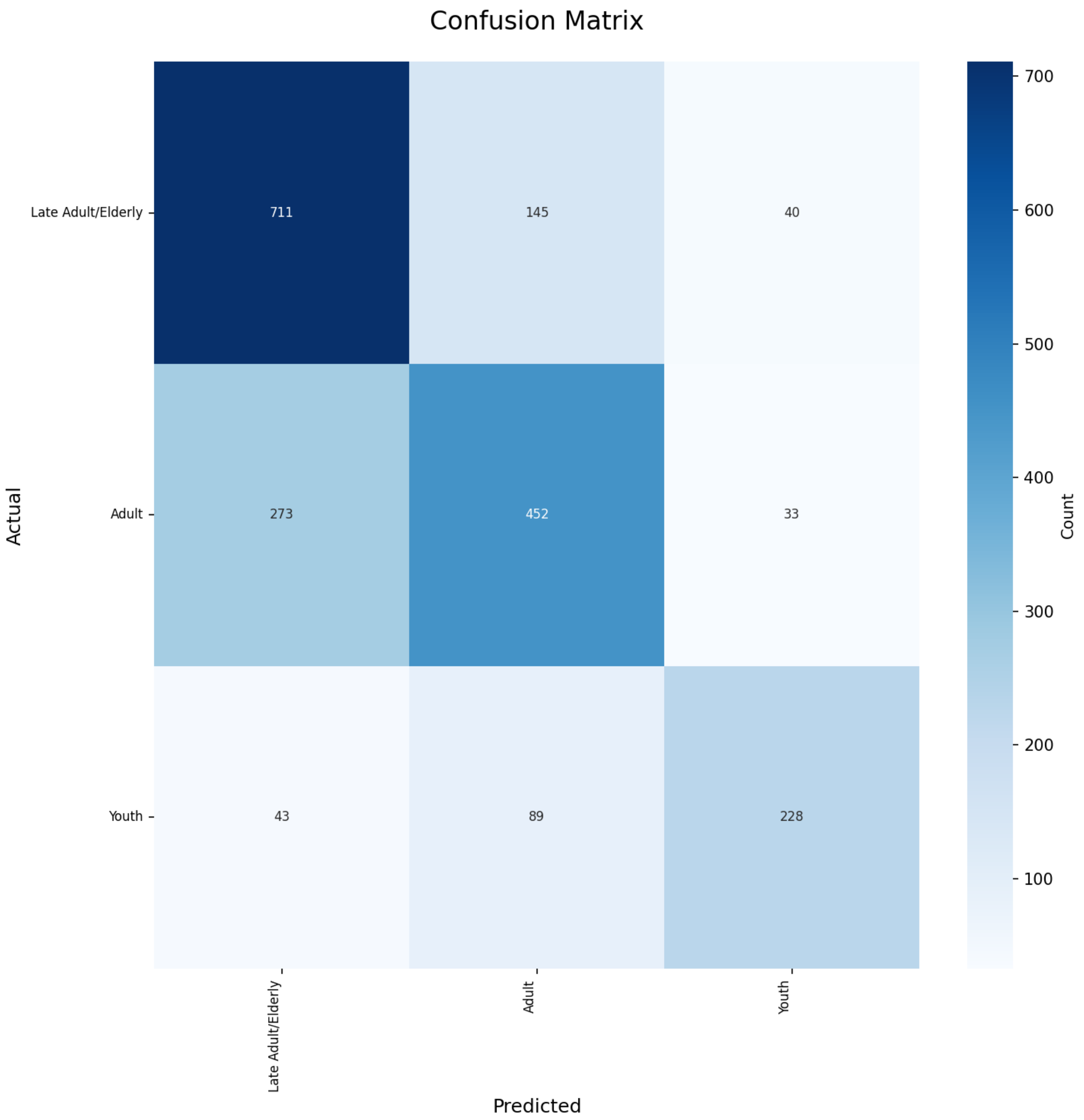


**Supplementary Figure S2.** The confusion matrix for age prediction result generated by finetuned CellPLM on 2024 holdout

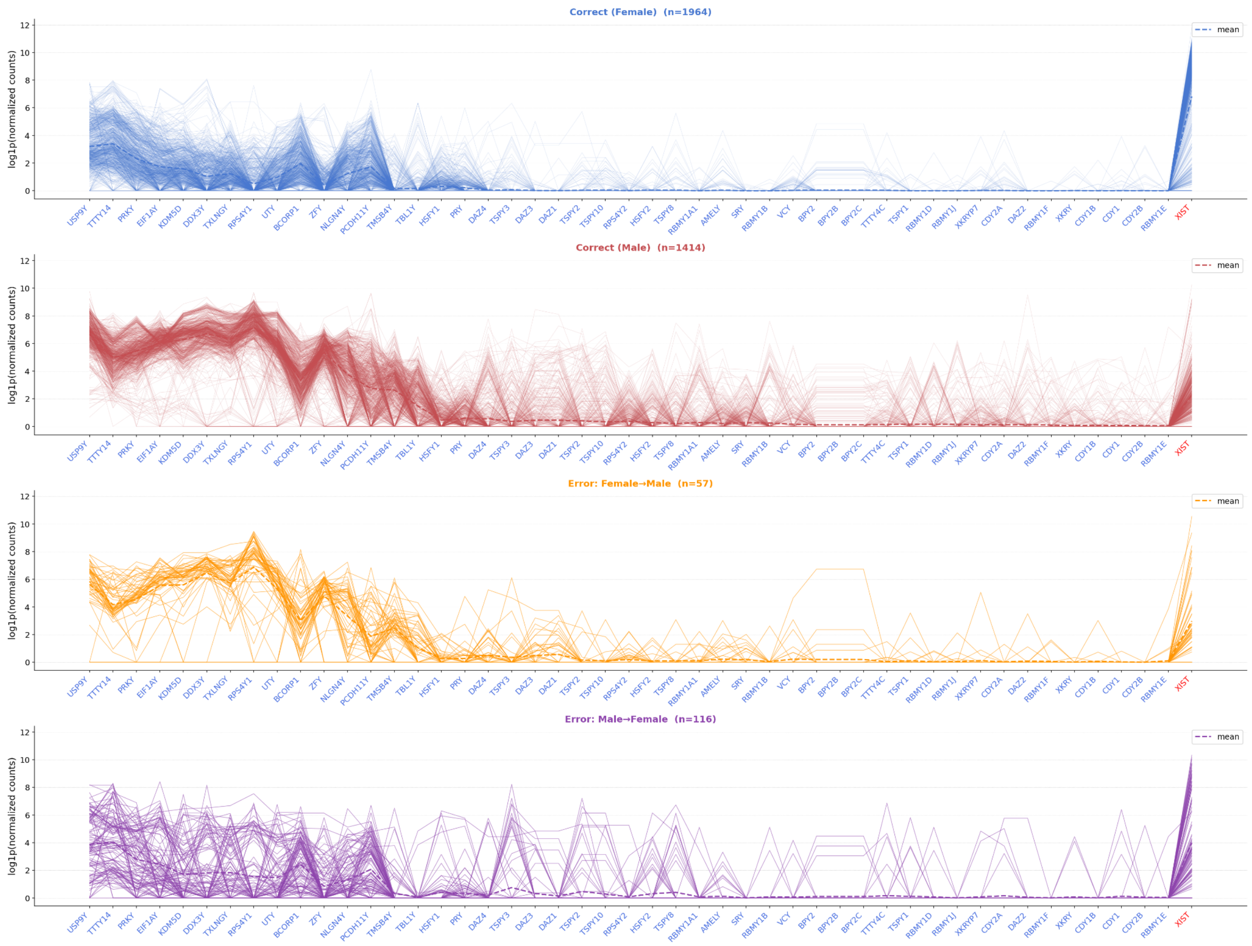


**Supplementary Figure S3. Sample-level expression profiles of sex-linked marker genes in the 2024 holdout cohort, stratified by classification outcome.** Each polyline represents one sample plotted across 48 sex-linked marker genes (log1p-normalized counts); dashed lines indicate group means. The four panels show correctly classified females (top left, n=1,964), correctly classified males (top right, *n* = 1,414), misclassified Female to Male samples (bottom left, *n* = 57), and misclassified Male to Female samples (bottom right, *n* = 116). X-axis tick labels show gene symbols; "XIST" (red) is the sole X-chromosome marker gene and is placed rightmost; all Y-chromosome gene labels are shown in blue.

**Supplementary Table S3**. Expert manual review confirmation rates for selected GEOMeta annotations in the prospective May 2026 GEO cohort

| **Reviewed field** | **Reviewed samples (*N*)** | **Reviewed GSEs (*N*)** | **Confirmed samples *n* (%)** | **Non-confirmed samples *n* (%)** |
|---|---|---|---|---|
| Disease | 1,140 | 11 | 1,140 (100.0%) | 0 (0.0%) |
| Tissue | 1,148 | 13 | 1,148 (100.0%) | 0 (0.0%) |
| RNA source | 1,148 | 13 | 741 (64.5%) | 407 (35.5%) |
| Age | 1,148 | 13 | 1,148 (100.0%) | 0 (0.0%) |
| Sex | 1,148 | 13 | 1,144 (99.7%) | 40 (0.003%) |
| Perturbation | 92 | 10 | 92 (100.0%) | 0 (0.0%) |
| Perturbation type | 92 | 10 | 92 (100.0%) | 0 (0.0%) |

**Supplementary Table S4:** Performance, runtime, and cost comparison of 5 Selected LLMs for GEOMeta annotation on perspective analysis

| Model | Runtime | | API cost ($) | | Final release | | CP release | |
|---|---|---|---|---|---|---|---|---|
| | Total (Hour) | Estimated per 6-sample study (min) | Total | Estimated per 6-sample study | GSM, *n (%)* | GSE, *n* | GSM, *n (%)* | GSE, *n* |
| **Gemini 3 Flash Preview** | 8.06 | 0.43 | 16.35 | 0.0146 | 6,292 (93.5%) | 221 | 883 (14.0%) | 42 |
| **Mistral Large 2512** | 8.51 | 0.46 | 13.20 | 0.0118 | 6,343 (94.2%) | 222 | 1,302 (20.5%) | 59 |
| **GPT-5 mini** | 9.32 | 0.50 | 12.04 | 0.0107 | 6,487 (96.3%) | 222 | 1,123 (17.3%) | 56 |
| **DeepSeek V4 Flash** | 13.80 | 0.74 | 2.75 | 0.0025 | 6,640 (98.6%) | 224 | 1,224 (18.4%) | 57 |
| **Qwen3.7 Plus** | 21.67 | 1.16 | 12.69 | 0.0113 | 6,516 (96.8%) | 223 | 1,262 (19.4%) | 59 |

**Supplementary Table S5.** Representative GEO age annotation edge cases

| GEO-reported info | Contextual interpretation | Final age |
|---|---|---|
| patient age: 58 | Human renal carcinoma donor sample | 58 Years |
| age: 50 | Study described infant blood collected at week 6-7 of life; the value therefore represented days rather than years | 50 Days |
| age: 14 weeks gestation; or Age: 137 days; or fetal liver week 17 | Study and tissue descriptions indicated fetal developmental timing | 14 Gestational Weeks; 137 Gestational Days; 17 Gestational Weeks |
| DOB: 18-Mar-1998; or sample date: 02-Feb-2010 | Age calculated from the reported date of birth and sample-collection date | 11 Years |
| CLiP_N, PY45, Female, 45 yr.; or NHDF IPSC from 44-year-old female | Age was reported in the sample title or source description rather than in a dedicated age field | 45 Years |
| age: 12 weeks during differentiation; 48 hpf; passage 75; mouse age; Age: 0; Age: 1964 | Experimental timing, passage number, non-human age, placeholder values, and implausible values were not treated as human donor age | NA |

**Supplementary Table S6.** Representative GEO tissue annotation examples

| Raw GEO term | Metadata context | Initial tissue extraction | Final standardized tissue |
|---|---|---|---|
| Lower segment myometrial tissue | Myometrial biopsy from uterus; the anatomical subregion was mapped to a broader tissue | Lower segment myometrium | Smooth muscle |
| Subcutaneous tumor | Xenograft from HCCLM3 liver cancer cells; the implantation site differed from the tumor tissue of origin | Subcutaneous tumor | Liver |
| Diencephalon | Fetal brain region mapping to a supported brain-region category | Diencephalon | Brain: Thalamus |
| Aorta-gonad-mesonephros | Embryonic hematopoietic developmental region without a supported tissue category | Aorta-gonad-mesonephros | NA |
| Adjacent normal tissue | Tumor-adjacent control from a liver carcinoma study; the underlying tissue was identified from study context | Adjacent normal liver tissue | Liver |
| PBMC | Isolated peripheral blood mononuclear cells; the cell preparation was mapped to its anatomical source | PBMC | Blood |

**Supplementary Table S7.** Full GEOMeta annotation schema and definitions for 27 metadata fields

| Field | Definition and annotation rule | Example output |
|---|---|---|
| GSM_ID* | Unique identifier of each sample (GSM) in GEO. | GSM8699331 |
| GSE_ID* | Series accession linking GSMs to the parent study. | GSE285257 |
| Seq_Type* | Sequencing type; one of BULK-RNA, SC-RNA, or Other. | BULK-RNA |
| Organism* | Scientific name of the source organism. | Homo sapiens |
| Strain | Specific strain or background information | NA |
| Genotype | Genetic constitution, standardized as “Knockout: [Gene]”, “Mutant: [Gene]”, etc. | NA |
| RNA_Library* | RNA library preparation method. | mRNA-based |
| RNA_Source* | Biological material from which RNA was extracted. | Cell Line: MOLM-13 |
| Tissue* | Anatomical tissue or organ associated with the sample, mapped to the released controlled tissue vocabulary. | Blood |
| Experimental_Setting* | Experimental context: In Vivo, Ex Vivo, or In Vitro. | In Vitro |
| Model_Type | Experimental model category (e.g., Cell Line, Organoid, PDX). | Cell Line |
| Disease* | Disease state associated with the sample, based on explicitly reported or contextually supported metadata. | Acute Myeloid Leukemia |
| GSE_Pert* | Whether the study included any perturbations (Yes/No). | Yes |
| GSM_Pert* | Perturbation status of the individual sample, represented as Perturbed, Control, or NA. | Perturbed |
| Perturbation* | Perturbation agent or intervention applied. | Decitabine |
| Pert_Dose* | Dose of the perturbation, with standardized units. | 100 nM |

| | | |
|---|---|---|
| Pert_Freq | Frequency of perturbation administration. | Every 24 Hours\| |
| Pert_Duration* | Total duration of perturbation exposure. | 72 Hours |
| Route_ Admin | Route of perturbation administration. | In Media |
| SampleType | Broad classification of the sample source. | Cell Line |
| SpecimenType | More specific description of biological material. | Cell Line |
| Race | Race reported for human donor samples; NA when unavailable or not applicable. | NA |
| Ethnicity | Ethnicity reported for human donor samples; NA when unavailable or not applicable. | NA |
| Age* | Standardized donor or developmental age with units; NA when unavailable or not applicable.ed, or not applicable. | 20 Years |
| Sex* | Donor sex from direct metadata or controlled tissue-based inference; NA when unavailable. | Male |
| Timepoint | Sampling time relative to treatment, event, or developmental stage. | Post-treatment Hour: 144 |
| Outcome | Treatment response, survival, or prognosis when reported. | Unknown |

* Included in the primary public GEOMeta release. Unmarked fields are part of the full annotation schema but are not included in the primary release.

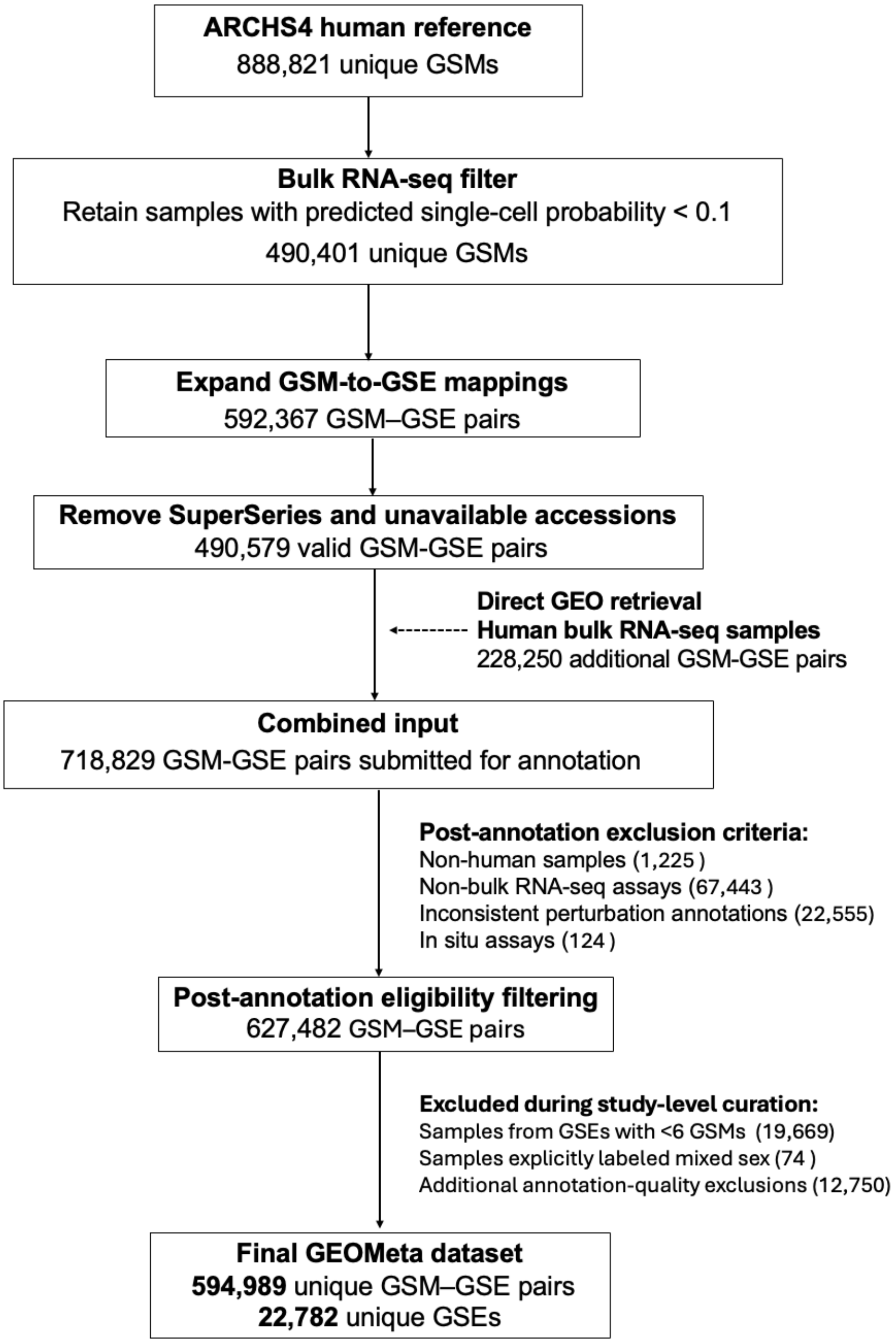


**Supplementary Figure S4.** Data retrieval, filtering and preprocessing workflow for the GEOMeta dataset

**Supplementary Table S8.** Annotation agents and output-field scope

| Annotation Agent | Output fields | Notes |
|---|---|---|
| Experimental context agent | Seq_Type, Organism, Strain, Genotype, RNA_Library, RNA_Source, Tissue, Experimental_Setting, Model_Type | Extracted jointly to improve consistency across related fields. |
| Biological context agent | Disease | Restricted to explicitly or contextually supported information. |
| Perturbation agent | GSE_Pert, GSM_Pert, Pert, Pert_Dose, Pert_Freq, Pert_Duration, Route_Admin | Preserved component order and attributes for combination perturbations. |
| Sample metadata agent | SampleType, Specimen_Type, Race, Ethnicity, Age, Sex, Timepoint, Outcome | Extracted jointly to reduce cross-field inconsistency. |

**Supplementary Table S9.** Age Group Definitions

| Age Group | Definition / Rule | Range (inclusive) |
|---|---|---|
| NA | No age or age category available or applicable (e.g., in vitro, cell lines, organoids, missing data). | |
| Infant | Fetus, neonate, or infant younger than 1 year old (includes terms like "Newborn", "0 Years"). | younger than 1 year |
| Pediatric | Early childhood to pre-adolescence. | 1 - 12.9 years |
| Adolescent | Teen years preceding adulthood. | 13 - 17.9 years |
| Adult | Metadata explicitly states "Adult," but no numeric age provided. | |
| Adults-20s | Young adults. | 18 - 29.9 years |
| Adults-30s | Adults in their thirties. | 30 - 39.9 years |
| Adults-40s | Adults in their forties. | 40 - 49.9 years |
| Adults-50s | Adults in their fifties to early sixties. | 50 - 64.9 years |
| Elderly | Metadata explicitly states "Elderly," but no numeric age provided. | |
| Elderly-1 | Early elderly. | 65 - 74.9 years |
| Elderly-2 | Mid-elderly. | 75 - 84.9 years |
| Elderly-3 | Late elderly | ≥ 85 years |

**Supplementary Table S10.** Summary of evaluation cohorts used in this study

| Evaluation cohort | Cohort size | Selection | Primary purpose | Evaluation strategy |
|---|---|---|---|---|
| Development benchmark | 1,000 samples | Manually selected during agent development to cover diverse experimental and metadata-reporting scenarios | Task-specific agent instruction development and internal evaluation | Agreement with manually reviewed annotations |
| Prompt-design comparison | 1,000 samples from 200 GSEs | Five randomly selected GSMs per GSE | Task-specialized versus generic Stage 1 prompt comparison | Annotation missingness and cross-prompt concordance |
| External-validation cohorts | DiSignAtlas: 19,879 matched samples; PharmGEO: 10,359 matched samples | GSM-level overlap after identifier harmonization | Disease, tissue and chemical-perturbation validation | Concordance with external curated resources |
| Representation-learning cohorts | 473,770 samples: 435,668 training; 24,496 study-level holdout; 13,606 2024 temporal holdout | Pre-2024 samples used for training, with 10% of pre-2024 GSEs retained as a study-level holdout; samples deposited in or after 2024 retained as a temporal holdout | Downstream model benchmarking | Label-prediction performance |
| May 2026 prospective cohort | 96 GSEs; 7,319 samples annotated; up to 1,148 samples reviewed | Size-stratified sample of human-only GSEs submitted in May 2026 | Prospective annotation and independent expert evaluation | Field-specific domain-expert confirmation in the reviewed subset |
| Internal multi-LLM benchmark | 118 samples from five GSEs | Studies submitted by our laboratory | Comparison and selection of 22 LLMs | Completeness, consensus concordance, runtime and cost |
| Dec 2025–Jan 2026 multi-LLM cohort | 225 GSEs; 6,733 GSM records annotated; 5,945 samples in the clean consensus release; 4,647 evaluated using CellPLM | December 2025–January 2026 GSEs with ≥ 6 ARCHS4-matched samples; parent SuperSeries excluded | Five-model prospective annotation and prospective CellPLM evaluation | Cross-model output completeness and consensus concordance; transcriptomic label-prediction performance |

**Supplementary Text 1.1**

For transparency and reuse, field-specific post-processing instructions are provided in the GEOMeta GitHub repository. Curated mapping resources are also provided for fields requiring controlled harmonization, including disease, tissue, RNA source, and chemical perturbation annotations. These files document the normalization rules, accepted standardized terms, and manually reviewed mappings used by the pipeline. GPT-5 was used for the post-processing tasks reported here.

**Supplementary Text 1.1:** Tissue Term Standardization and Controlled-Vocabulary Mapping

To achieve consistent tissue annotations across studies, raw tissue and cell source descriptions from GEO were standardized through a two-stage process consisting of rule-guided normalization followed by controlled-vocabulary mapping and review.

In the first stage, raw free-text annotations were normalized using a standardization agent with explicit predefined rules. The tissue standardization agent received explicit instructions to apply predefined, rule-guided transformations. These rules enforced abbreviation expansion, synonym consolidation, controlled handling of anatomical subregions, and consistent formatting, producing intermediate standardized terms suitable for controlled-vocabulary mapping. Core normalization rules included standardized capitalization and formatting; abbreviation expansion; consolidation of synonymous or related anatomical terms; controlled reduction of anatomical subregions to parent tissues; and assignment of NA when no biologically valid mapping could be established. Representative transformations included “iWAT” to “Adipose tissue”, “Lymph nodes” to “Lymphoid tissue”, “Heart left ventricle” to “Heart”, and “Melanoma” to “Skin”.

In the second stage, normalized terms were mapped to a controlled vocabulary derived from the Human Protein Atlas (HPA) Tissue Atlas. Standardized tissue terms were mapped, when supported, to one of 37 top-level tissue categories: Eye, Retina, Heart, Skeletal muscle, Smooth muscle, Adrenal gland, Parathyroid gland, Thyroid gland, Pituitary gland, Lung, Bone marrow, Lymphoid tissue, Liver, Gallbladder, Testis,

Epididymis, Prostate, Seminal vesicle, Adipose tissue, Brain, Choroid plexus, Salivary gland, Esophagus, Tongue, Stomach, Intestine, Pancreas, Kidney, Urinary bladder, Breast, Vagina, Cervix, Endometrium, Fallopian tube, Ovary, Placenta, and Skin. Brain-derived samples were assigned either Brain or one of 13 supported region-specific labels using the format Brain: <Region>, including Brain: Cerebral cortex, Brain: Cerebellum, Brain: Basal ganglia, Brain: Thalamus, Brain: Hypothalamus, Brain: Midbrain, Brain: White matter, Brain: Amygdala, Brain: Choroid plexus, Brain: Pons, Brain: Medulla oblongata, Brain: Hippocampal formation, and Brain: Spinal cord. When brain origin was clear but the specific region could not be determined, the sample was assigned Brain.To improve coverage beyond the core HPA ontology, 7 additional categories were introduced for underrepresented or anatomically specialized sources, including Blood, Nasal, Nasopharynx, Oropharynx, Umbilical Vein, head and neck, and Synovium regions. When datasets described cultured or isolated cell populations, tissue of origin was inferred from sample metadata or lineage information and mapped to the most appropriate canonical category. Samples with ambiguous, conflicting, or malformed anatomical descriptors were excluded from harmonization to preserve ontological integrity.

All final standardized tissue mappings were reviewed manually for biological plausibility and cross-field consistency. Reviewed mappings were retained for reuse in subsequent annotation runs.

**Supplementary Text 1.2** Age Annotation Standardization and Age-Group Derivation

Age annotations were retained only when explicitly and unambiguously reported in the GEO metadata. Values were standardized to numeric plus unit format (e.g., “25 Years,” “8 Months”) with uniform capitalization and spacing. Implausible or malformed values were removed.  In Stage 2, an age-group derivation agent performed a controlled inference task to assign the Age_Group annotation from the standardized age values using predefined developmental bins (Supplementary Table S9). Samples labeled only as “Adult” or “Elderly” without numeric values were retained under their respective broader categories. In vitro systems (cell lines, organoids) and other cases where biological age was not applicable were assigned “NA”.

**Supplementary Text 1.3:** Disease Term Curation, Standardization, and Ontology Mapping

Disease annotations were harmonized using a multi-stage workflow that integrates deterministic rules, LLM-assisted standardization, ontology-based retrieval, and manual quality control. The goal was to generate ontology-consistent, biologically interpretable disease labels while retaining provenance and rare but meaningful disease concepts.

**Step 1. Initial Disease Annotation**

Disease labels were first generated by the biological context annotation agent using both GSE-level and GSM-level metadata.. Each sample was assigned to one of four principal categories:

1) Normal: Healthy or non-diseased control samples with explicit evidence of normal physiological state.

2) Adjacent Normal: Histologically normal tissue derived from a donor with a documented disease condition.

3) Disease-Associated: Samples with explicit disease names or well-established disease origins (such as MDA-MB-231 cells being associated with breast cancer.).

4) No Disease Mentioned: Samples lacking reliable disease information, including most iPSC-derived systems and purely methodological or modeling datasets.

Disease names were extracted directly when explicitly stated in GEO metadata. Inference was performed only when contextual evidence was strong and unambiguous. Samples containing conflicting, vague, or non-resolvable disease descriptions were assigned “NA” and excluded from downstream disease analyses.

**Step 2. Disease Term Standardization**

Raw disease annotations were standardized using a field-specific disease standardization agent task . Standardization included abbreviation expansion, such as

SLE to Systemic Lupus Erythematosus; capitalization normalization; synonym consolidation; and removal of non-essential modifiers while preserving core disease identity.

Following agent-based normalization, additional rule-based filtering and manual review were applied. Terms representing anatomical locations, procedures, molecular alterations, experimental conditions, or non-disease descriptors were removed. Samples associated with multiple unrelated diseases or ambiguous disease contexts were excluded from downstream ontology mapping. The resulting standardized disease terms were used as inputs for controlled-vocabulary alignment.

**Step 3. Frequency Consolidation and Term Filtering**

A frequency table of standardized disease terms was generated to identify unique entries for ontology alignment. Rare disease entities absent from CTD MEDIC but supported by biologically valid terminology and recurring across multiple samples, such as STAT2 Deficiency and X-linked Reticulate Pigmentary Disorder, were retained under their standardized names. Ambiguous, or non-disease concepts were removed at this stage.

**Step 4. Mapping to CTD MEDIC**

Standardized disease terms were mapped to the Comparative Toxicogenomics Database (CTD) MEDIC vocabulary, which integrates MeSH and OMIM disease concepts. The mapping pipeline consisted of four coordinated layers:

(a) Deterministic Matching
Disease terms first underwent normalization procedures including case standardization, punctuation removal, and synonym normalization. Exact and synonym matching was then performed against CTD MEDIC disease names and synonym tables. Specific disease subtypes could also be mapped to established parent disease concepts when supported, such as Triple-Negative Breast Cancer to Breast Neoplasms.

(b) Similarity-Based Candidate Retrieval

Terms without deterministic matches were evaluated using TF–IDF similarity, fuzzy string matching, and token overlap scoring. The highest-ranked CTD MEDIC candidates were retrieved programmatically and supplied to the disease mapping agent for constrained evaluation.

(c) Constrained Mapping-Agent Selection

The disease mapping agent evaluated the standardized disease term and the retrieved CTD MEDIC candidates. High-confidence deterministic matches were used directly, while ambiguous or low-similarity cases underwent contextual evaluation against retrieved ontology candidates.

The mapping instructions required the agent to:

- prioritizing canonical parent disease concepts over highly specific molecular or numbered subtypes,
- normalizing cancer terms to established neoplasm classes,
- excluding overly broad or non-specific descriptors,
- rejecting composite multi-disease phrases,
- and avoiding unsupported ontology assignments when no reliable CTD concept existed.

Each accepted mapping returned the canonical CTD disease term together with associated ontology identifiers and mapping rationale.

(d) Manual Review and Quality Control

Mappings with low similarity scores (below 0.75) or unresolved ontology assignments were flagged for manual review. Additional curation removed non-disease concepts, overly broad pathological descriptions, and unstable multi-condition annotations. Rare diseases were retained only when biologically meaningful and supported across sufficient samples (≥10 samples). Unresolvable cases were assigned “NA.” Final reviewed mappings were saved in reusable mapping tables for subsequent annotation runs.

**Step 5. Final Disease Annotation Output**

Each curated disease annotation includes the following ontology-aligned fields:

- Canonical CTD MEDIC disease name
- MeSH, OMIM, and DOID identifiers
- Alternative identifiers, definitions, and synonyms
- hierarchical tree-number information; and
- high-level disease-category mappings.

Rare disease terms retained outside CTD MEDIC were preserved under their standardized names. This curation strategy yields precise, ontology-consistent disease annotations while preserving rare but biologically meaningful disease entities outside standard vocabularies.

**Supplementary Text 1.4:** Perturbation-Related Fields Standardization and Controlled Inference

Perturbation-related annotations were harmonized through a structured workflow to ensure consistent representation of perturbation status, perturbation names, dose, duration, frequency, and route of administration across studies. This process was applied to all perturbation types and served as the basis for downstream chemical-specific mapping.

**Step 1. Standardization of perturbation names and attributes**

Raw perturbation terms were normalized using field-specific standardization agent and rule-based post-processing. The core perturbation name was separated from non-identifying modifiers such as dose, duration, concentration, frequency, and route of administration, while these attributes were retained in their corresponding structured fields. Standardization also addressed naming variation by expanding abbreviations, consolidating synonyms, and enforcing consistent capitalization, hyphenation, and formatting. Chemical aliases, brand names, cytokines, antibodies, genetic perturbation descriptions, and environmental conditions were standardized to consistent representations where supported by the metadata.

Combination perturbations were preserved as combined terms using consistent separators, such as “+” for co-treatments and “;” for distinct genetic modifications. This preserved perturbation ordering and treatment context. Control-related terms were standardized separately to distinguish untreated, vehicle, solvent, and placebo conditions from active perturbations. Genetic perturbations were normalized using consistent descriptors such as Knockout, Knockdown, Overexpression, Mutant, or Transgenic, while environmental perturbations were simplified to standardized descriptive terms such as hypoxia, diet, radiation, or stress.

**Step 2. Controlled inference of perturbation type**

A perturbation-type derivation agent performed a controlled inference task to derive the perturbation type from the standardized perturbation names using predefined output categories. These included CTL for control or untreated conditions, CP for chemical perturbation, BIO for biological agents, KO for knockout, KD for knockdown, OE for overexpression, ES for environmental stress, VIR for viral infection, OTHER for deliberate perturbations not captured by the main categories, and NA when no intentional perturbation was supported.

Inference rules were based on the semantic content of the standardized perturbation name, with explicit handling of common biological agents, genetic modifications, environmental exposures, and control conditions. For combination perturbations, category codes were assigned in the same order as the corresponding perturbation agents to preserve experimental structure. The distinction between OTHER and NA was maintained to separate deliberate but unclassified perturbations from samples without evidence of intentional intervention.

**Step 3. Dose and duration standardization outcomes**

At the vocabulary level, standardization reduced perturbation-dose annotations from 3,975 distinct cleaned values to 1,162 release values and perturbation-duration annotations from 1,143 to 433 values. After excluding the release categories NA and Others, 1,160 standardized dose values and 431 standardized duration values remained.

At the sample level, perturbation dosage was reported for 114,094 samples (19.2% of the release). After applying the standardization rules, which retained only single-dose values with approved concentration or mass units, 67.9% of samples with reported dose information were retained as standardized dose values, whereas 32.1% were classified as “Others.” Perturbation duration was reported for 26.5% of the release, of which 95.4% of samples with reported duration information were retained as standardized single-duration values and 4.6% were classified as “Others.”

Together, these steps converted heterogeneous perturbation descriptions into standardized perturbation labels and structured perturbation-type annotations while preserving key experimental context. This harmonization supports identification of treated and control samples and facilitates downstream analysis of chemical, genetic, biological, and environmental perturbations across studies.

**Supplementary Text 1.5:** Chemical Perturbation Standardization and PubChem Mapping

Chemical perturbations were curated using a dedicated multi-step workflow designed to ensure accurate compound identification, standardized naming, and interoperability with external chemical resources. This workflow builds on the general perturbation standardization described in Supplementary Text 1.4 and was applied exclusively to samples annotated as chemical perturbations.

**Step 1. Chemical perturbation name standardization**

All 39,775 raw perturbation terms were first normalized using the general perturbation standardization workflow to 31,860 unique standardized names. Variant expressions of the same compound were collapsed through frequency-based aggregation to generate a standardized set of chemical perturbation names.

**Step 2. Identification of chemical perturbation terms**

Perturbation types were inferred from standardized perturbation names using the predefined perturbation classification scheme described in Supplementary Text 1.4. Terms labeled as chemical perturbations included both single-compound entries and composite chemical perturbations, such as terms joined by "+". In total, 20.8% (6,640) unique perturbation terms were assigned to chemical or composite chemical perturbation categories.

**Step 3. Restriction to single-compound chemical perturbations**

To avoid ambiguous compound-level assignments, PubChem and structural mapping were restricted to single-compound chemical perturbation terms. Composite perturbations involving multiple compounds were excluded. This filtering retained 4,721 single-compound chemical terms for downstream mapping. Of the 4,721 eligible single-compound terms, 66.7% were successfully mapped to PubChem, representing 2,787 unique PubChem compounds and 72,117 sample-level chemical perturbation records.

**Step 4. PubChem mapping and match classification**

Single-compound chemical perturbation terms were mapped to PubChem compound records. Mapping prioritized exact title matches and synonym matches, with additional support for abbreviation expansion and close-match review when needed. PubChem results were accepted only when the returned compound name or synonym supported the queried perturbation term. Ambiguous or unverified results were retained for review rather than assigned a compound record. Each mapped term was assigned a final mapped compound name (Matched PubChem Name), PubChem CID, SMILES strings, mapping category, and PubChem URL where available.

**Step 5. Structural annotation**

For mapped compounds, structural descriptors were retrieved from PubChem. CanonicalSMILES values were extracted for compounds with valid PubChem CIDs. Canonical SMILES strings were available for 99.99% of the 72,117 mapped sample-level records. These descriptors support downstream applications such as drug signature analysis, cheminformatics, and compound representation learning.

**Supplementary Text 1.6:** RNA Source Standardization and Mapping

RNA source annotations were standardized to represent the biological material from which RNA was extracted, including tissues, isolated cell populations, established cell lines, biofluids, and other source materials. Raw RNA source values were first normalized using field-specific standardization agent to reduce variation in capitalization, pluralization, abbreviations, cell-marker formatting, and redundant descriptors. The post-processing step preserved original term order and applied source-specific standardization rules rather than introducing unsupported labels.

The reviewed mapping resource for RNA source was constructed by applying the predefined RNA-source standardization and mapping rules, followed by manual review and correction. These rules defined the allowed final label formats and the handling of anatomical tissues, isolated cell populations, cell lines, biofluids, organoids, generic non-informative labels, blood-derived materials, tumor-related terms, and marker-defined immune-cell subsets. After review, accepted mappings were incorporated into the mapping file and used for subsequent annotation runs.

For any new RNA source terms not already present in the reviewed mapping file, an RNA-source mapping agent applied the same predefined rules used to construct the reviewed mappings. Final mapped labels were restricted to Tissue: xx, Cells: xx, Cell Line: xx, Biofluid: xx, Other, or null. Null was used only when the source term was intentionally non-informative or not suitable for release-level source assignment.

The mapping rules distinguished anatomical tissues, isolated cell populations, cell lines, organoids, biofluids, and generic non-informative labels. Clear anatomical terms were mapped to controlled tissue-style labels when supported by the metadata context. Adjacent-normal or disease-status labels were mapped to the underlying anatomical source only when the source was explicit, such as adjacent normal cervix or adjacent non-tumor liver tissue. Generic labels such as tumor, biopsy, extracellular vesicles, or organoid were not treated as sufficiently specific RNA source labels unless additional anatomical or cellular context was available. Organoid-containing terms were not automatically mapped to the tissue of origin unless the mapping was already reviewed or explicitly supported.

Cell-line terms were additionally mapped against a DepMap cell-line model metadata reference (obtained from the Broad Institute DepMap Data Portal). Candidate terms were normalized and compared with canonical cell-line names and curated aliases in the reference table. When a confident reference match or manually reviewed alias match was identified, the corresponding canonical cell-line label was assigned. Recurrent unmatched terms were retained only when they occurred in at least 30 samples; less frequent unmatched terms were collapsed to Cell Line: Other to avoid introducing unstable, sparsely represented labels.

Low-confidence, ambiguous, or unsupported RNA source terms were flagged for review rather than treated as fully curated mappings. This design separates reviewed reusable mappings from provisional automated mappings and allows newly observed source terms to be evaluated and incorporated into the mapping resource in future annotation runs. The final release files include the mapped RNA source field, while internal audit outputs retain mapping status, mapping method, confidence, reasoning, and review-required indicators for traceability.

RNA-source standardization reduced 17,112 submitter-derived terms to 953 standardized labels, representing a 94.4% reduction in label diversity. Of the 592,201 samples with non-missing RNA-source annotations, 97.8% were assigned a standardized RNA-source label using the reviewed mapping resource.

### Supplementary Text 2. External validation methods

**Supplementary Text 2.1** DiSignAtlas preprocessing, disease and tissue concordance assessment

Disease and tissue annotations in GEOMeta were evaluated using DiSignAtlas, a manually curated resource of disease-associated transcriptomic signatures with harmonized metadata. DiSignAtlas provides standardized disease and tissue annotations, as well as sample groupings used in disease signature comparisons. We first identified overlapping GEO studies (GSEs) between DiSignAtlas and GEOMeta. DiSignAtlas records were expanded to the sample level by parsing GSM identifiers from case_accession and control_accession fields, so that each row corresponds to a (GSE_ID, GSM_ID) pair.GSM identifiers were parsed from the reported case_accession and control_accession fields, and the GEO series accession was treated as the GSE identifier. DiSignAtlas metadata are curated at a study-signature level and indexed by an internal curation identifier (dsaid). During sample-level expansion, multiple DiSignAtlas records were occasionally associated with the same *(GSE_ID, GSM_ID)* pair, reflecting alternative disease interpretations curated under different dsaid entries. Because GEOMeta does not contain dsaid information, sample pairs associated with multiple DiSignAtlas records were excluded before integration. Only unique sample-level mappings were retained for downstream analyses.

Disease annotations from GEO metadata were compared against DiSignAtlas disease annotations using a hierarchical matching framework. Samples annotated as *Normal, No Disease Mentioned, Adjacent Normal*, or *Healthy* were excluded prior to disease concordance analysis. Agreement was classified into four categories: *Direct match* (identical disease terms), *Synonym match* (equivalent disease concepts based on ontology synonyms), *Generalized match* (biologically equivalent or closely related disease concepts ,e.g., subtype versus parent disease, determined through expert manual review), and Unmatched (no concordance). This allowed differentiation between strict term agreement and biologically meaningful agreement.

To further assess disease annotation consistency, GEO-derived disease labels were compared with DiSignAtlas case/control assignments. In DiSignAtlas, case/control labels indicate a sample's role in a disease signature comparison, where *Case* represents the disease or condition of interest and *Control* represents the reference group. GEOMeta samples annotated as Normal, Adjacent Normal, or No Disease Mentioned were grouped as non-disease, and all other samples were classified as disease-associated. Concordance was defined as non-disease samples labeled as *Control* and disease-associated samples labeled as *Case*. This comparison was performed at the GSM level across all matched samples.

For tissue evaluation, GEOMeta tissue annotations were compared with DiSignAtlas tissue labels at the GSM level. Concordance was first assessed across all matched samples. Samples with missing DiSignAtlas tissue annotations were retained and analyzed separately to distinguish annotation absence from true disagreement. Tissue labels were standardized using the same controlled vocabulary applied to GEOMeta tissue region fields, based on mappings derived from the Human Protein Atlas. Primary concordance analyses were performed using standardized labels to avoid conflating formatting differences with annotation disagreement. To further examine tissue disagreements, we restricted the analysis to samples with non-missing tissue labels in both resources. Raw and standardized tissue labels were compared separately to distinguish formatting or synonym differences from persistent discrepancies.

**Supplementary Text 2.2** PharmGEO preprocessing and chemical perturbation concordance assessment

Perturbation annotations in GEOMeta were externally validated using PharmGEO, a curated resource of drug-response transcriptomic studies from GEO. PharmGEO annotations are provided at the contrast level, where each record specifies treated and control sample groups within a study, rather than at the individual sample level. To enable direct comparison, PharmGEO records were expanded to sample-level entries by parsing treated and control sample identifiers for each contrast. Control samples were labeled as controls without assigned drug identifiers, while treated samples were assigned standardized drug names curated by PharmGEO. For studies with multiple treatment arms or combination therapies, all associated drugs were retained as composite annotations to preserve experimental context. The resulting sample-level dataset was restricted to human samples and merged with GEOMeta annotations using exact matching on study and sample identifiers.

Perturbation concordance was evaluated through four comparisons: (1) agreement between GEOMeta perturbation labels and PharmGEO standardized compound names; (2) consistency between PharmGEO treated/control assignments and GEOMeta perturbation type; (3) consistency between PharmGEO treated/control assignments and GEOMeta sample-level perturbation status; and (4) internal consistency within GEOMeta between perturbation type and sample-level perturbation status. Binary concordance indicators were assigned to each comparison.

For compound-level comparison, an initial exact string-matching approach was applied after normalization of compound names. To further characterize and resolve apparent disagreements, a frequency table was generated for all pairs of GEOMeta perturbation labels and PharmGEO standardized drug names among samples initially classified as discordant. This table was used to systematically identify recurrent mismatch patterns, including chemical synonyms, abbreviated versus full compound names, salt or formulation variants, and cases where one resource represented a subset of a multi-drug perturbation. Based on this analysis, a curated equivalence table was constructed and applied to re-evaluate drug agreement. Binary concordance flags were assigned before and after reconciliation to quantify the impact of this refinement.
